# A photoactivable natural product with broad antiviral activity against enveloped viruses including highly pathogenic coronaviruses

**DOI:** 10.1101/2021.07.09.451770

**Authors:** Thomas Meunier, Lowiese Desmarets, Simon Bordage, Moussa Bamba, Kévin Hervouet, Yves Rouillé, Nathan François, Marion Decossas, Fézan Honora Tra Bi, Olivier Lambert, Jean Dubuisson, Sandrine Belouzard, Sevser Sahpaz, Karin Séron

**Affiliations:** Univ Lille, CNRS, INSERM, CHU Lille, Institut Pasteur de Lille, U1019-UMR 9017-CIIL-Center for Infection and Immunity of Lille, Lille, France; Univ Lille, Université de Liège, Université de Picardie Jules Verne, JUNIA, UMRT 1158 BioEcoAgro, Métabolites spécialisés d’origine végétale, F-59000 Lille, France; UFR Sciences de la Nature, Université Nangui Abrogoua, BP 801 Abidjan 02, Côte d’Ivoire; Univ Bordeaux, CBMN UMR 5248, Bordeaux INP, F-33600 Pessac, France

## Abstract

The SARS-CoV-2 outbreak has highlighted the need for broad-spectrum antivirals against coronaviruses (CoVs). Here, pheophorbide a (Pba) was identified as a highly active antiviral molecule against HCoV-229E after bioguided fractionation of plant extracts. The antiviral activity of Pba was subsequently shown for SARS-CoV-2 and MERS-CoV, and its mechanism of action was further assessed, showing that Pba is an inhibitor of coronavirus entry by directly targeting the viral particle. Interestingly, the antiviral activity of Pba depends on light exposure, and Pba was shown to inhibit virus-cell fusion by stiffening the viral membrane as demonstrated by cryo-electron microscopy. Moreover, Pba was shown to be broadly active against several other enveloped viruses, and reduced SARS-CoV-2 and MERS-CoV replication in primary human bronchial epithelial cells. Pba is the first described natural antiviral against SARS-CoV-2 with direct photosensitive virucidal activity that holds potential for COVID-19 therapy or disinfection of SARS-CoV-2 contaminated surfaces.

## Introduction

The COVID-19 pandemic has highlighted the lack of specific antiviral compounds available against coronaviruses (CoVs). COVID-19 is caused by the severe acute respiratory syndrome coronavirus 2 (SARS-CoV-2), the third identified human CoV causing severe pneumonia (*1*–*3*). Before 2003, coronaviruses were known to cause severe diseases in animals, but human CoVs, such as HCoV-229E and OC-43, were mainly associated with common colds, and only rarely with severe outcomes (*4*). The severe acute respiratory syndrome coronavirus (SARS-CoV) outbreak in 2003 was the first emergence of a highly pathogenic human CoV. The second highly pathogenic coronavirus, identified in 2012 in Saudi Arabia, is the Middle-East respiratory syndrome coronavirus (MERS-CoV), which is still endemically present up to date. SARS-CoV-2 is highly related to SARS-CoV (*5*). The tremendous efforts of the scientific community worldwide to counteract the COVID-19 pandemic have rapidly led to the development of highly efficient vaccines that are now administered worldwide. Unfortunately, the emergence of SARS-CoV-2 variants in different regions of the world might render the vaccines less efficient and may necessitate a booster vaccination every year which might not be achievable for billions of human beings in all parts of the world. Therefore, to get rid of this pandemic and face future epidemics, it is assumed that not only vaccination is necessary but also the availability of efficient antiviral treatments. Before the emergence of SARS-CoV-2, no specific antiviral was commercially available for treatment of CoV infections. Due to the urgent need for antivirals against SARS-CoV-2, many researchers have focused their investigations on repurposing available drugs. Unfortunately, until now, none of them has been able to significantly reduce severe outcomes in patients. High content screening *in vitro* of approved drugs identified some potential interesting antiviral molecules that have still to be tested in clinic on patients with COVID-19 (*6*–*8*). Protease and polymerase inhibitors are also widely investigated in *in vitro* studies (*9*). Recently, White *et al*. identified plitidepsin, an inhibitor of the host protein eEF1A as a potential antiviral agent for SARS-CoV-2 (*10*). To date, only synthetic neutralizing monoclonal antibodies have been approved for emergency usage in newly infected patients.

Coronavirus are members of the *Coronaviridae* family within the order *Nidovirales*. CoVs are enveloped viruses with a positive single-stranded RNA genome of around 30 kb. The genome encodes 4 structural proteins, the nucleocapsid (N), the spike (S), the envelope (E), and the membrane (M) proteins. The S protein is important for the interaction of the viral particle with the cellular host receptor, being angiotensin-converting enzyme 2 (ACE2) for SARS-CoVs (*11*, *12*), dipeptidyl peptidase 4 (DPP4) for MERS-CoV (*13*) and aminopeptidase N (APN) for HCoV-229E (*14*). Once attached, the virus releases its genome into the cytosol by fusion of the viral envelope with a host membrane. This fusion process is mediated by the S protein, a class I fusion protein, and can occur either at the plasma- or at the endosomal membrane. Viral class I fusion proteins are typically synthesized as inactive precursor proteins and require proteolytical activation by cellular proteases to acquire their fusion competent state. The host-cell protease TMPRSS2 has been shown to be necessary for plasma membrane fusion of many coronaviruses including SARS-CoV-2 (*12*, *15*, *16*), whereas cathepsins are often involved in fusion processes at endosomal membranes (*17*).

It is estimated that about 80% of the global population rely on traditional medicine to treat infectious diseases. Plants are a natural source of compounds with a structural diversity that is much higher than those obtained by chemical synthesis. Many of these compounds have proven their antiviral activity *in vitro*. To date, some reports describe the antiviral activity of natural compounds on coronavirus including SARS-CoV-2, but many of them are *in silico* analyses without any *in vitro* or *in vivo* evidence (*18*, *19*).

Here we show that pheophorbide a (Pba) isolated from *Mallotus oppositifolius* (Geiseler) Müll.Arg. (Mo, *Euphorbiaceae)* leave crude extract after bioguided fractionation has antiviral activity against various CoVs, including HCoV-229E, MERS-CoV and SARS-CoV-2, but also against other enveloped viruses, such as yellow fever virus (YFV), hepatitis C virus (HCV) and Sindbis virus (SINV). Moreover, we demonstrate that Pba is an antiviral photosensitizer directly acting on the viral particle, thereby impairing the virus-cell fusion step.

## Results

### Pheophorbide a (Pba) isolated from the crude methanolic extract from *Mallotus oppositifolius* (Geiseler) Müll.Arg. is highly active against HCoV-229E

Fifteen plants methanolic extracts from the Ivorian pharmacopeia, which were initially screened for their anti-HCV activity (Bamba *et al.*, submitted), were tested against HCoV-229E-Luc, a luciferase recombinant version of HCoV-229E, which allows for a rapid and easy screening of diverse molecules *in vitro*. Seven of the fifteen extracts significantly reduced HCoV-229E infection (Fig. 1A), whereas none of the extracts showed cytotoxicity in Huh-7 cells at the tested concentrations (Bamba *et al.*, submitted). The *Mallotus oppositifolius* (Mo) crude extract was the most active and therefore selected for further analyses. A bioguided fractionation was performed to find out the active compound(s) in this plant. This revealed that the methylene chloride (MC) partition was the most active on HCoV-229E (Fig. 1B) and was therefore chosen for fractionation by centrifugal partition chromatography (CPC), leading to 10 fractions (F1-F10). Among these fractions, F7 was the most active and subjected to further fractionation by preparative high performance liquid chromatography (HPLC), leading to 9 subfractions (F7.1-F7.9). F7.7 was the most active of them and seemed to contain only one molecule. This dark green product was analysed by LC-UV-MS, which revealed only one peak (m+H = 593,3) with two maxima of absorption in the visible light (409 and 663 nm). This information was used for a Dictionary of Natural Product search (https://dnp.chemnetbase.com/), indicating that this molecule could be pheophorbide a (Pba). F7.7 purity and chemical structure was further confirmed by nuclear magnetic resonance (NMR)(data not shown).

**Fig. 1.**
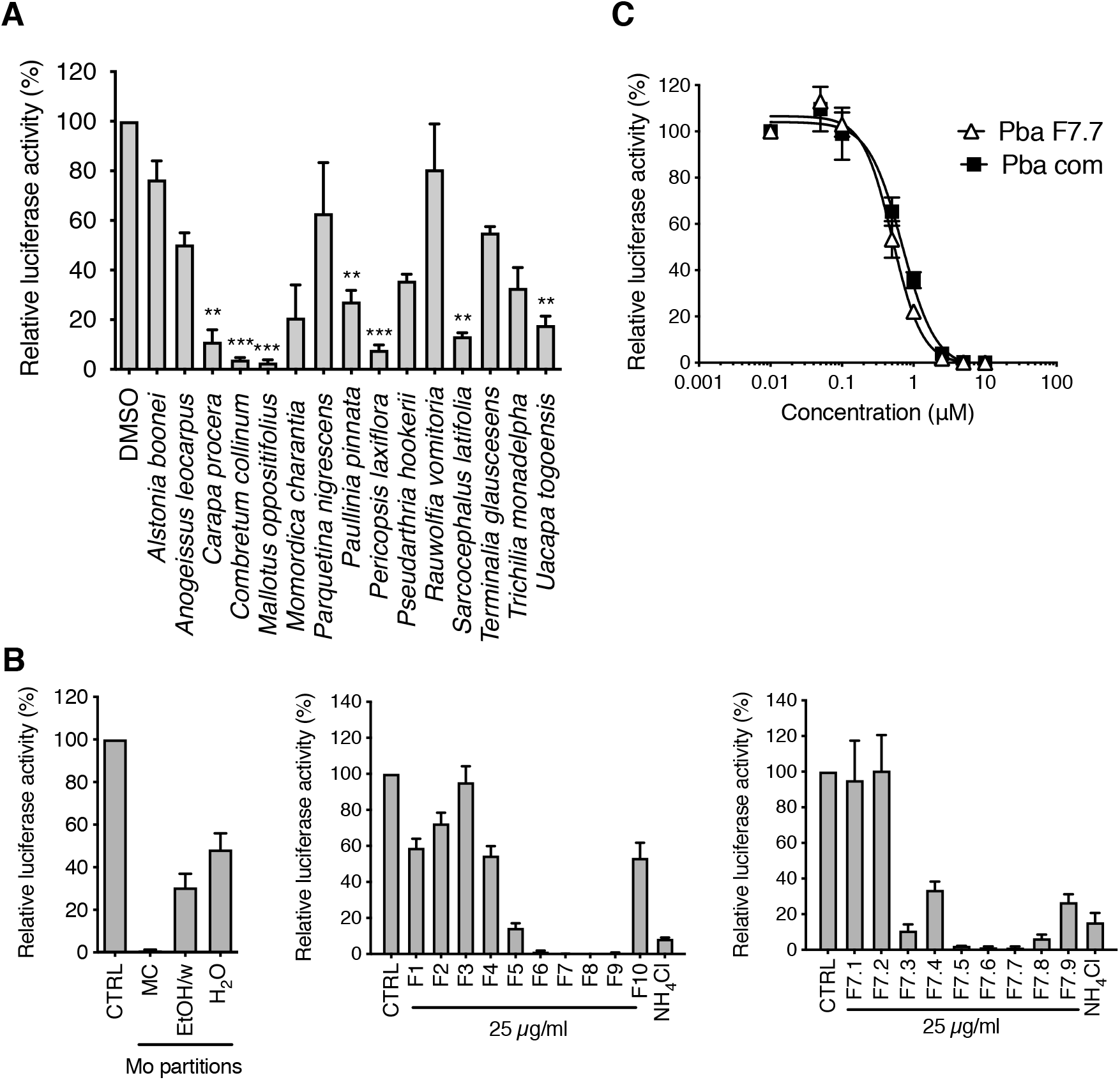
Identification of Pba as active compound in *Mallotus oppositifolius* using bioguided fractionation of plant extracts. (**A**) Huh-7 cells were inoculated with HCoV-229E in the presence of various plant extracts at 25 μg/mL. Cells were lysed 7 h post-inoculation and luciferase activity quantified. (**B**) Huh-7 cells were inoculated with HCoV-229E in the presence of sub-extracts of Mo (methylene chloride, MC; ethanol/water (50:50), EtOH/w; water, H_2_0; left), fractions of Mo MC sub-extract (middle), or sub-fractions of F7 fraction (right panel) at 25 μg/mL. Cells were lysed 7 h post-inoculation and luciferase activity quantified. (**C**) Huh-7 cells were inoculated with HCoV-229E-Luc in the presence of Pba extracted from Mo (Pba F7.7) or commercial Pba (Pba com) at different concentrations. At 1 h p.i., cells were washed and fresh compounds were added to the cells for 6 h after which cells were lysed to quantify luciferase activity. Data are expressed relative to the control DMSO. Results are expressed as mean ± SEM of 3 experiments.

To confirm that Pba was the active compound of the Mo extract, a dose-response experiment was performed with both the pure compound isolated in F7.7 and commercial Pba against HCoV-229E-Luc in Huh-7 cells. As shown in Fig. 1C, IC_50_ values were comparable for both natural and commercial Pba (0.51 μM and 0.54 μM, respectively), confirming that Pba was indeed the active substance of Mo.

### Pba is active against several human CoVs at non-cytotoxic concentrations

Prior to the assessment of its broad-spectrum activity against various CoVs, the cytotoxicity of Pba was determined in different cells lines. As Pba is known to be a photosensitizer it was assumed that light could affect its toxicity. MTS assays were performed in similar conditions than the infection procedures. Cells were incubated with Pba at different concentrations and taken out of the incubator after 1 h to change the medium and left for 10 min under light exposure in the biosafety cabinet (BSC), after which they were replaced in the incubator for 23 h. In parallel, plates were kept for 24 h in the dark. Pba did not exhibit any toxicity at concentrations up to 120 μM when left in the incubator for 24 h without light exposure, whereas it showed some toxicity when the cells were shortly exposed to light, with CC_50_ values of 4.4 ± 1.2 μM, 5.8 ± 1.9 μM and 5.5 ± 2.0 μM for Huh-7, Vero-E6 cells, and Vero-81, respectively (Fig. S1). Taken together, these results show that the cytotoxicity of Pba in cell culture depends on light exposure.

The antiviral activity of Pba was tested on different human CoVs. It was first confirmed on HCoV-229E by measuring infectious titers (Fig. 2) with an IC_50_ value of 0.1 μM, resulting in a selectivity index of 44. Similarly, infection inhibition assays were performed with various non-cytotoxic concentrations of Pba against SARS-CoV-2 and MERS-CoV. As shown in Fig. 2, Pba had a strong antiviral effect against both highly pathogenic coronaviruses, with IC_50_ values of 0.18 μM for both SARS-CoV-2 and MERS-CoV, resulting in a selectivity index of 32 and 24, respectively.

**Fig. 2.**
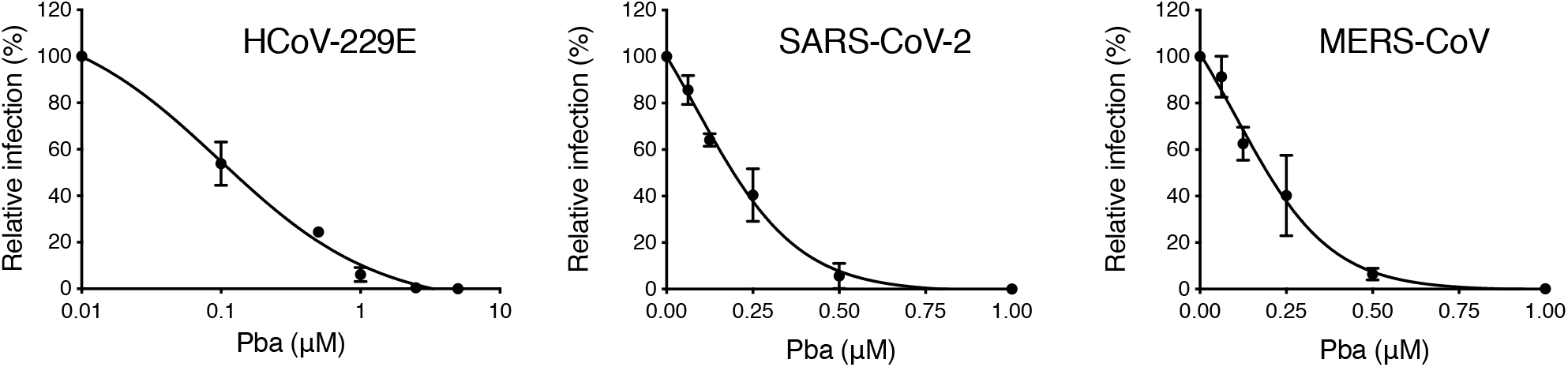
Pba inhibits various HCoVs. Cells were inoculated with HCoV-229E (Huh-7 cells), SARS-CoV-2 (Vero-E6 cells) and MERS-CoV (Huh-7 cells) in presence of various concentrations of Pba. At 1 h p.i, cells were washed and fresh compounds were added to the cells for 9 h (HCoV-229E) or 16 h (SARS-CoV-2 and MERS-CoV) and the supernatants were collected for infectivity titration. Results are expressed as mean ± SEM of 3 experiments.

### Pba is an inhibitor of coronavirus entry by direct action on the particle

Coronaviruses fusion is triggered by proteolytic cleavage of the spike protein. Depending on cellular proteases available, fusion can occur after endocytosis of the virus or directly at the cell surface. It has been demonstrated that HCoV-229E and SARS-CoV-2 fusion at the plasma membrane depends on the expression of the TMPRSS2 protease (*12*, *20*), and for many coronaviruses the entry via fusion at the plasma membrane has been shown to be the most relevant pathway *in vivo* (*21*). To determine if Pba was able to inhibit both entry pathways, its antiviral activity was tested in Huh-7, and Huh-7-TMPRSS2 cells, the latter being obtained after transduction with a TMPRSS2 lentiviral expression vector. No difference in antiviral activity of Pba against HCoV-229E infection was observed in the presence or absence of TMPRSS2 (Fig. 3A). Similar results were obtained with SARS-CoV-2 in Vero cells (our unpublished observation). These results show that Pba exhibits antiviral activity whatever the entry pathway used.

**Fig. 3.**
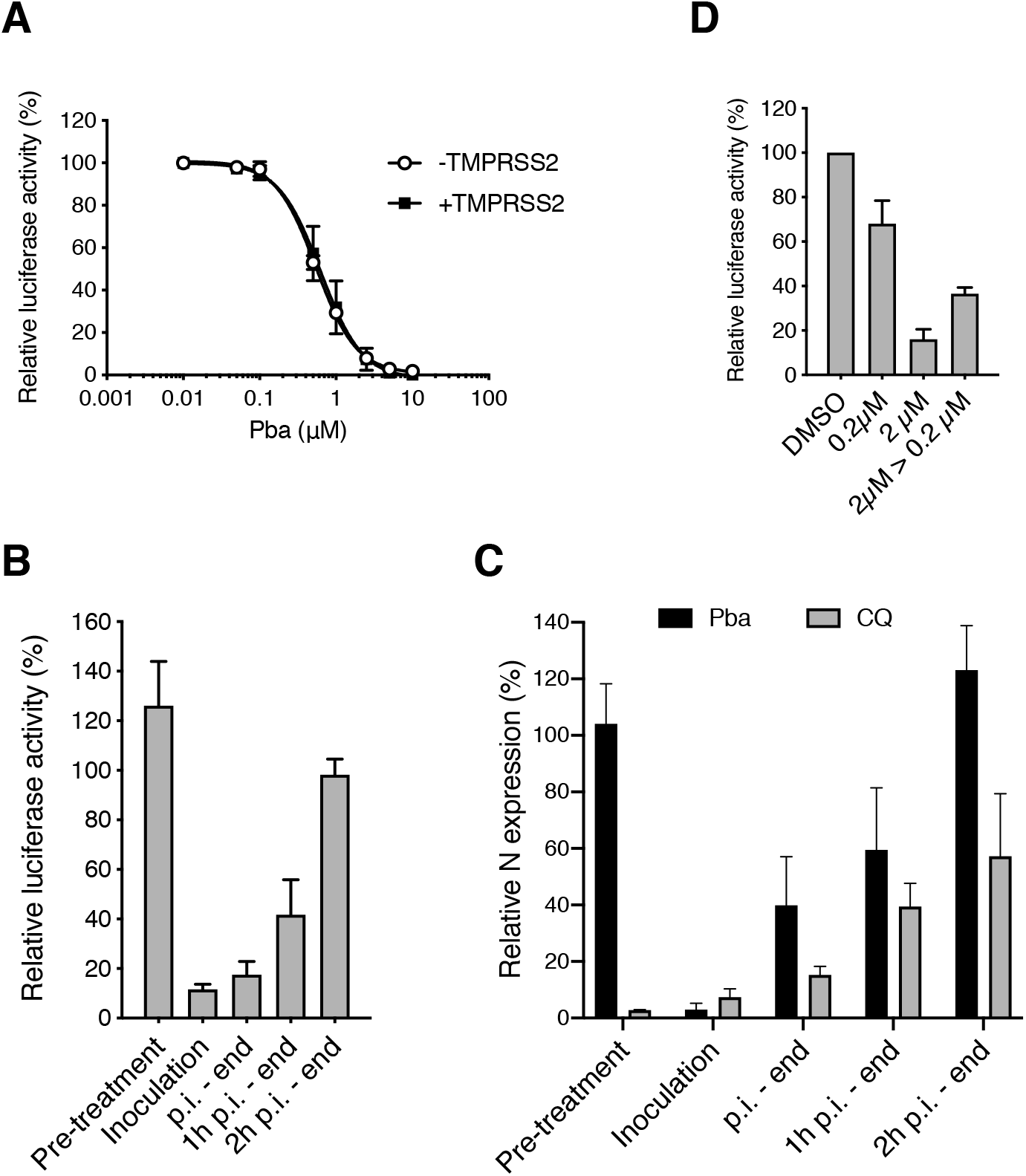
Pba inhibits viral entry by a direct action on the viral particle. (**A**) Huh-7 and Huh-7-TMPRSS2 cells were inoculated with HCoV-229E-Luc in the presence of various concentrations Pba. At 1 h p.i, cells were washed and fresh compounds were added to the cells for 6 h after which the cells were lysed to quantify luciferase activity. Data are expressed relative to the control DMSO. (**B**) Pba at 2 μM was added at different time points during infection of Huh-7-TMPRSS2 cells by HCoV-229E-Luc, either 1 h before inoculation (pre-treatment), or 1 h during inoculation (Inoculation), or for 6 h post-inoculation (p.i.-end), after 1 h post-inoculation till the end (1 h p.i.-end), or 2 h post-inoculation till the end (2 h p.i.-end). Cells were lysed 7 h after the inoculation and luciferase activity quantified. (**C**) A similar experiment was performed in Vero-81 cells inoculated with SARS-CoV-2 in the presence of Pba at 1 μM or chloroquine at 10 μM at different time points. Cells were lysed 16 h post-inoculation and the viral nucleocapsid protein was detected by Western blot. The graph represents the quantification of the band intensity corresponding to the N protein, relative to the DMSO control for each time point. (**D**) Huh-7 cells were inoculated with HCoV-229E-Luc in the presence of 0.2 or 2 μM Pba, or with HCoV-229E-Luc previously treated with 2 μM Pba and then diluted 10 times, leading to a concentration of 0.2 μM Pba for the inoculation period (2 μM > 0.2 μM). The amount of virus used for inoculation was kept constant in the different conditions and all the samples were exposed to the light for 10 min. At 7 h post-inoculation, cells were lysed and luciferase activity was quantified.

To gain insights into the mechanism of action of Pba, a time-of-addition assay with Pba was performed during HCoV-229E or SARS-CoV-2 infection. Therefore, Pba was added at different time points before, during or after inoculation. All experiments were performed under BSC’s light exposure. For both viruses, no inhibition of infection was observed when Pba was added to the cells before inoculation (pre-treatment of the cells), whereas a strong inhibition of infection was noticed when Pba was present during the virus inoculation step (Fig. 3B and 3C). When Pba was added only after inoculation, its inhibitory effect rapidly dropped and normal levels of infectivity were observed again when Pba was added more than 2 h after inoculation. Chloroquine is a well-known inhibitor of SARS-CoV-2 replication *in vitro*, as it inhibits virus-cell fusion after endocytosis by preventing endosomal acidification. As shown in Fig. 3C, the Pba and chloroquine inhibition curves were rather similar, showing that Pba could inhibit virus entry. It is worth noting that normal levels of infectivity were observed when Pba was added more than 2 h after inoculation, whereas chloroquine-treated cells remained at around 60% of normal levels which may be due to inhibition of the second round of entry since our experimental conditions are compatible with reinfection. Taken together, these results suggest that Pba is inhibiting virus entry.

Since no antiviral activity was observed with cells treated with Pba prior to inoculation, we wondered whether Pba targets the virus instead of the cells. To test this hypothesis, HCoV-229E-Luc was pre-incubated for 30 min at a concentration 10 times higher than that used during inoculation. For this experiment, HCoV-229E-Luc was pre-incubated with Pba at 2 μM for 30 min, then diluted to reach a concentration of Pba of 0.2 μM for inoculation, a concentration which does not severely impact HCoV-229E-Luc infection as shown above. In parallel, cells were directly inoculated with HCoV-229E-Luc at 0.2 and 2 μM as a control. The results clearly show that when HCoV-229E-Luc was pre-treated with Pba at high concentration (2 μM) before inoculation at low concentration (0.2 μM), the antiviral activity was much stronger than when inoculation was performed in the presence of 0.2 μM Pba without any pre-treatment (Fig. 3D). Taken together, these results indicate that Pba inhibits HCoV entry by a direct effect on the viral particle.

### Pba is an inhibitor of viral fusion

Virus entry can be divided into two different steps, firstly the viral attachment to the cell surface, and secondly the fusion of the virus envelope with cellular membranes. To further define the mode of action of Pba, experiments were performed with HCoV-229E, which can be manipulated in a lower containment facility. To determine a potential effect at the attachment step, Huh-7-TMPRSS2 cells were incubated with HCoV-229E in the presence or absence of Pba at 4°C for 1 h. These conditions block endocytosis but allow virus attachment to the cell surface. Cells were rinsed with PBS and the amount of virions attached to the surface was determined by quantification of viral genomes by qRT-PCR. As shown in Fig. 4A, only a slight and non-significant decrease in RNA levels was observed in the presence of Pba, indicating that Pba barely affect virus attachment. The action of Pba on the fusion step was investigated in virus-cell fusion assay by using trypsin as an exogenous protease to induce coronavirus membrane fusion at the plasma membrane (*22*, *23*). Cells were treated with NH_4_Cl to inhibit fusion in the endocytic pathway and viruses were bound at the cell surface at 4°C. Then, fusion was induced by a short trypsin treatment at 37°C. Entry at the cell surface was more efficient in presence of trypsin compared to the control (Fig. 4B), which is consistent with other reports (*22*, *23*). In addition, Pba at both 0.5 and 1 μM strongly inhibited infection levels in trypsin-mediated fusion conditions in a similar range compared to the inhibition seen with the control. Taken together, these results indicate that Pba inhibits entry at the fusion step and not by preventing attachment.

**Fig. 4.**
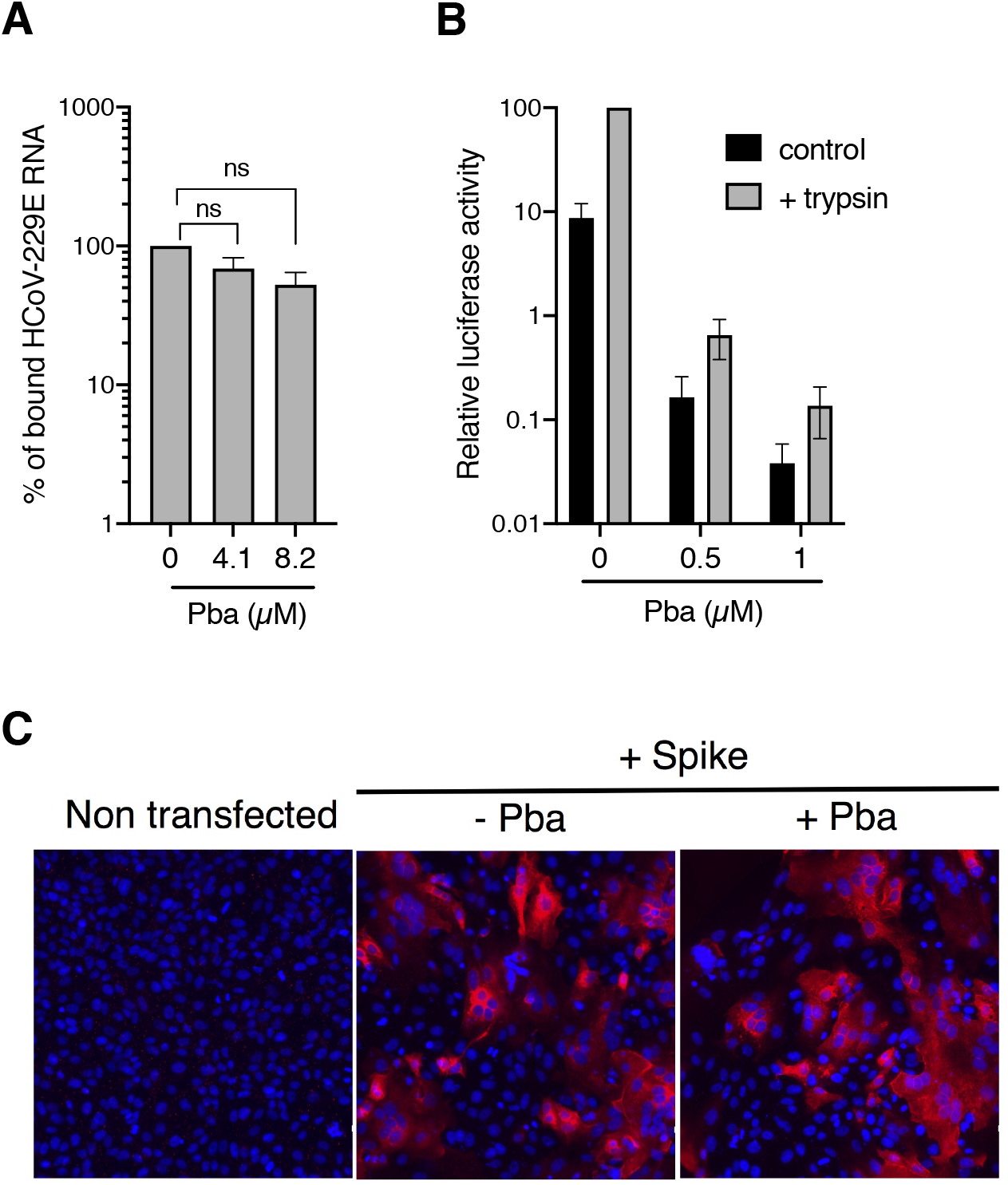
Pba inhibits viral entry at the fusion step. (**A**) Huh-7-TMPRSS2 cells were inoculated with HCoV-229E for 1 h at 4**°**C in the presence of DMSO, or 4.1 and 8.2 μM Pba. Cells were washed thrice with ice-cold PBS, and total RNA was extracted. Bound HCoV-229E virions were detected by quantification of HCoV-229E gRNA by qRT-PCR. Relative binding is expressed as the percentage of the control (DMSO) for which the 100% value was arbitrarily attributed. Mean values ± SEM (error bars) of three independent experiments are presented. n.s., not significant. (**B**) HCoV-229E was incubated with Pba at different concentration and was bound to Huh-7 cells for 1 h in the absence (control) or presence (trypsin) of NH_4_Cl at 4°C. In the later condition, fusion was induced by 3 μg/mL trypsin for 5 min at 37 °C in the presence of NH_4_Cl. Cells were lysed 7 h post-infection and luciferase activity quantified. Infectivity is expressed as the percentage of the control (DMSO) for which the 100% value was arbitrarily attributed. Mean values ± SEM (error bars) of three independent experiments are presented. (**C**) Vero-81 cells transiently expressing SARS-CoV-2 spike protein were incubated with or without Pba at 1 μM from 6 to 24 h p.i., after which syncytia were visualized by immunofluorescence. Images were acquired on an Evos M5000 imaging system (Thermo Fisher Scientific).

Two viral structures are involved in the virus-cell fusion, namely the spike protein and the viral membrane itself. To find out if Pba targets the spike, a cell-cell fusion assay was performed. For this, the SARS-CoV-2 spike protein was transiently expressed by plasmid transfection in Vero-81 cells. At 6 h post-transfection, the medium was replaced with medium containing either DMSO or 1 μM Pba until 24 h p.t.. As shown in Fig. 4C, spike induced cell-cell fusion (apparent as syncytium formation) occurred equally well in control (DMSO)- and Pba-treated conditions, indicating that Pba cannot prevent cell-cell fusion induced by the viral spike protein. Taken together these results are not in favour of an effect of Pba on the viral spike fusion protein.

### The antiviral activity of Pba is light-dependent and targets the viral membrane

Given that Pba is photoactivable, we wondered whether its antiviral effect is light-dependent. We therefore inoculated Huh-7 cells with HCoV-229E in the presence of Pba under different light exposure conditions. As shown in Fig. S2, the antiviral activity of Pba is light-dependent. A typical feature of photosensitizers is that the concentration required for its biological properties decreases upon increase of time of exposure to the light. To see whether Pba behaves as other photosensitizers, HCoV-229E was pre-incubated with Pba at various concentrations (0.02, 0.2 and 2 μM) and exposed to the normal white light of the laminar flow cabinet for different durations (ranging from 5 to 80 min). As shown in Fig. 5A, results clearly showed that with a same concentration of Pba, increased inhibitory effect could be observed with longer light exposure times.

**Fig. 5.**
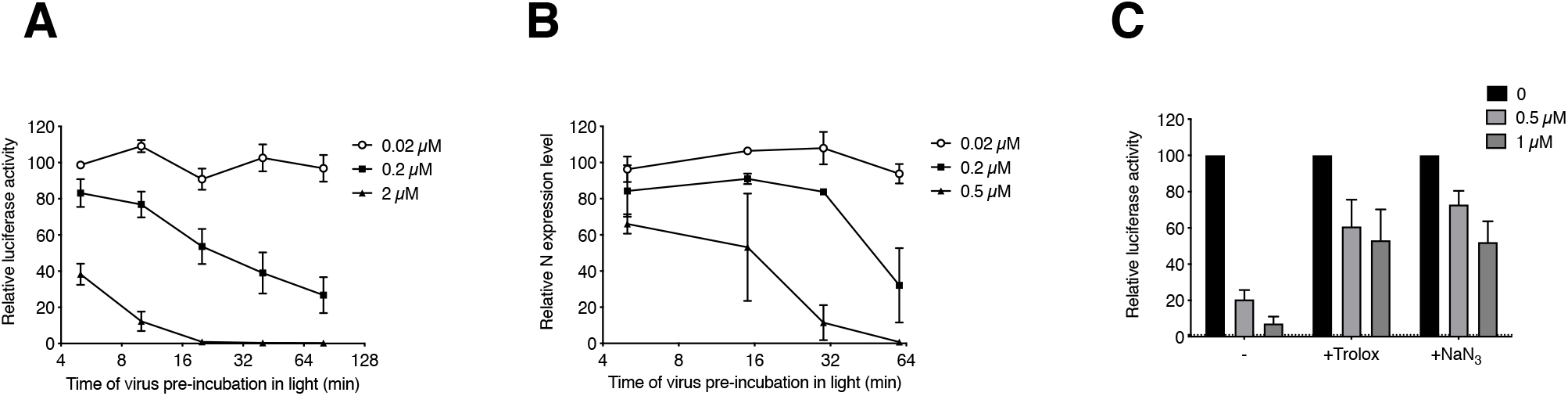
The antiviral activity of Pba depends on light exposure and its ability to generate singlet oxygen species. HCoV-229E-Luc (**A**) and SARS-CoV-2 (**B**) were incubated with Pba at given concentration under the light of the laminar flow cabinet. At different time points of light exposure, the mixture was used to inoculate Huh-7 cells or Vero-81 cells, respectively. At 7 h (HCoV-229E-Luc) or 16 h (SARS-CoV-2) post-inoculation, cells were lysed and infection quantified as described previously. (**C**) Pba at 0.5 or 1 μM was mixed with 10 mM Trolox or 10 mM NaN_3_ in DMEM and HCoV-229E-Luc was added to the mixture prior to inoculation of Huh-7 cells for 1 h at 37°C. Inoculum and compounds were removed and replaced with culture medium for 6 h. Cells were lysed and luciferase activity quantified. Infectivity is expressed as the percentage of infection relative to the control (DMSO) to which the 100% value was arbitrarily attributed. Mean values ± SEM (error bars) of three independent experiments are presented.

Similar results were observed with SARS-CoV-2 (Fig. 5B). Together, these data indicate that the anti-coronavirus properties of Pba depend on its dynamic photoactivation.

Photodynamic inactivation (PDI) of microorganisms typically results from the onset of reactive oxygen species (ROS), including free radicals or singlet oxygen species (^1^O_2_), generated when the light-activated photosensitizer falls back to its ground state, thereby transferring its energy to molecular oxygen (resulting in the onset of ^1^O_2_) or initiating photochemical reactions with ROS generation. Two mechanisms of activation have been described, either type I reactions in which the photosensitizer activates a substrate that generates reactive oxygen species (ROS), or type II reactions in which the photosensitizer directly generates singlet oxygen (^1^O_2_). Subsequently, these species can damage various micro-organism structures, such as nucleic acids, proteins or lipids (*24*, *25*). To determine if the antiviral activity of Pba also depends on ROS or ^1^O_2_ generation, infection was performed in the presence of quenchers that are able to trap these generated oxygen species. Two ^1^O_2_ quenchers were used, a water-soluble analogue of vitamin E, Trolox, and NaN_3_ (Fig. 5C). HCoV-229E was mixed with the quenchers Trolox and NaN_3_ both at 10 mM, and Pba was added at the inoculation step. Then the cells were rinsed and fresh culture medium was added for 6 h. The results clearly showed that both Trolox and NaN_3_ were able to reduce the action of Pba (Fig. 5C), indicating that the antiviral activity of Pba is mediated by the generation of ^1^O_2_ after photoactivation.

Vigant *et al.* clearly demonstrated that the generation of ^1^O_2_ by the lipophilic photosensitiser LJ001 induces the phosphorylation of unsaturated phospholipid of viral membranes, and changes the biophysical properties of viral membranes, thereby affecting membrane fluidity and/or increasing rigidity (*24*). As a result, the change of fluidity and/or rigidity of the viral membrane impairs its ability to undergo virus-cell fusion. We therefore wondered whether a similar action of the photosensitizer on the lipids of the viral envelope might also explain our observation that Pba is able to inhibit HCoV-229E fusion by targeting the viral particle. We hypothesized that the Pba-induced membrane rigidity may render the virus less sensitive to a shrinkage effect induced by an osmotic stress. Therefore, HCoV-229E was incubated with Pba either in the dark or under light exposure for 30 min, and subjected to osmotic shock with 400 mM NaCl before fixation with 4% PFA. Fixed viral particles were visualized by cryo-electron microscopy (CryoEM). As shown in Fig. 6, intact virions with their characteristic spikes at the surface can be observed in untreated conditions (control). The addition of Pba either in the dark or under light condition did not affect the overall morphology of virions in normal medium conditions. Interestingly, when intact virions were subjected to an osmotic stress by increasing NaCl concentration from 100 mM to a final concentration of 400 mM, the virions shape was altered due to a shrinkage of the viral membrane. When virions incubated with Pba in the dark were subjected to osmotic shock, similar alteration of viral shape was observed. However, in the presence of Pba and under light condition, no membrane deformation was observed suggesting that the virus was more resistant to osmotic shock. These results clearly show that light-activated Pba modify the mechanical properties of the viral envelope by increasing its stiffness. This Pba-induced increase in the envelope rigidity likely prevents the membrane deformation needed to undergo virus-cell fusion. Knowing that there might be an effect of the light, the spike-induced cell-cell fusion assay as shown in Fig. 4C was repeated with short light exposure of the spike-transfected cells every 2 h after addition of Pba. Even with regular exposure to the light, no effect of Pba was seen on the spike-mediated cell-cell fusion (data not shown), further excluding that Pba additionally targets the spike protein.

**Fig. 6.**
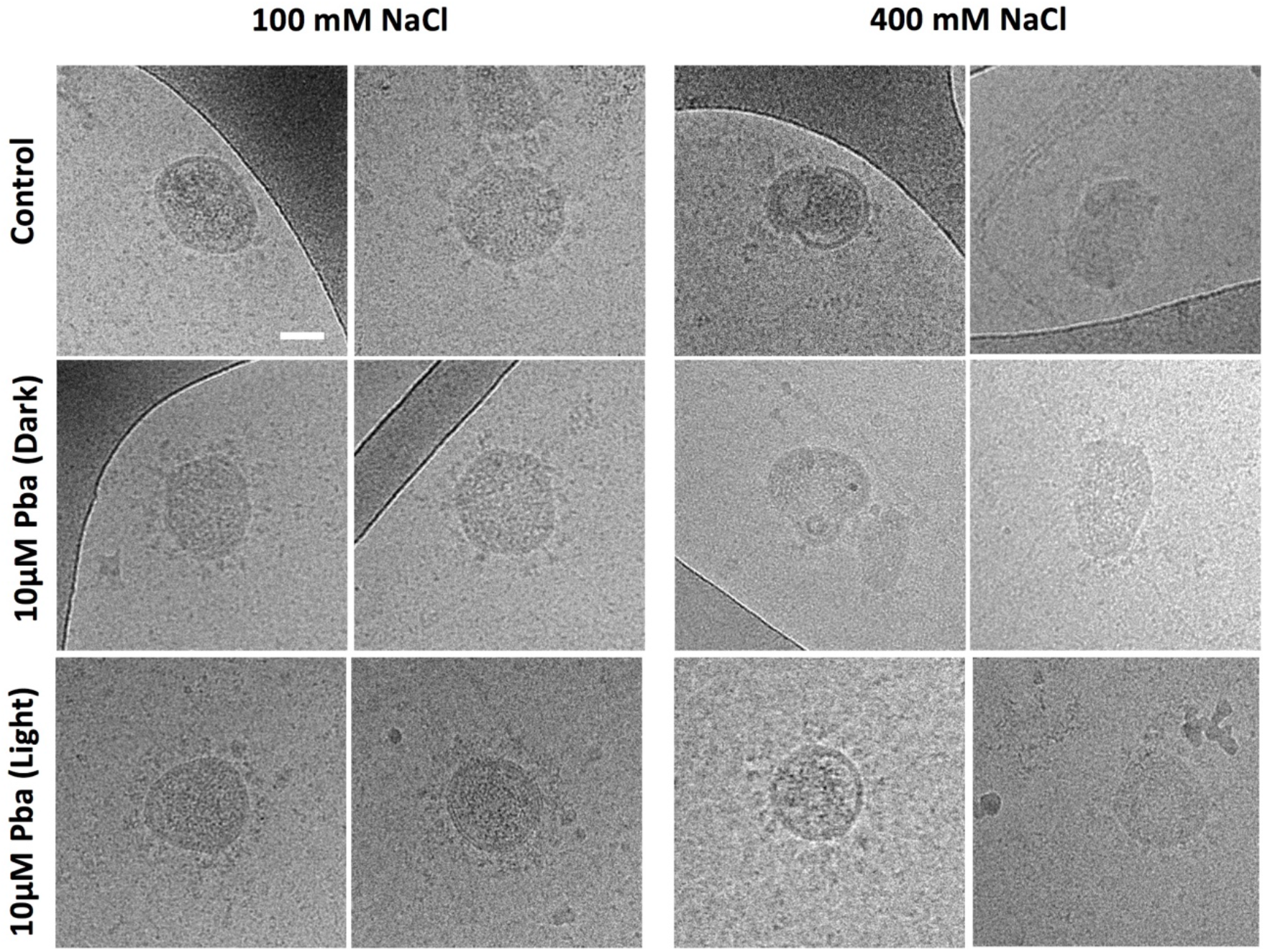
Pba renders virions resistant to osmotic shock. HCoV-229E were incubated in the presence or absence of Pba at 10 μM with or without 30 min light exposure, after which the particles were subjected to normal medium conditions (100 mM NaCl) or to an osmotic shock of 400 mM NaCl for 30 sec. Virions were fixed with PFA and samples were treated for CryoEM observation. Images are representative of 30 independent images of 2 independent experiments. Scale bar: 50 nm.

### Pba is a broad-spectrum antiviral that targets viral membranes of several enveloped viruses

In contrast to viral proteins, viral membranes are derived from the host cell and hence are not virus specific. This suggests that Pba might have broader antiviral activity against enveloped viruses. Indeed, antiviral activities against other viruses have already been reported for Pba or related molecules. These include HCV, HIV (Human immunodeficiency virus), and IAV (Influenza A virus) (*26*–*28*). To confirm the broad-spectrum activity of Pba, its antiviral activity was tested on pseudotyped viral particles with envelope proteins of VSV (vesicular stomatitis virus), HCV, SARS-CoV-2, and MERS-CoV. Pseudoparticles were pre-treated with 0.5, 1 or 2 μM Pba and exposed or not to the light for 30 min prior to inoculation. The results clearly showed that Pba inhibited pseudoparticle infection regardless of the nature of the viral envelope protein, but only under light conditions (Fig. 7A). Next, the antiviral activity of Pba was tested for coxsackievirus (CVB4, non-enveloped), Sindbis virus (SINV, enveloped), hepatitis C virus (HCV, enveloped), and yellow fever virus (YFV, enveloped). Whereas Pba was not active on the non-enveloped virus CVB4, it showed a clear antiviral effect against the 3 enveloped viruses HCV, YFV and SINV (Fig. 7B). Taken together, these results confirm the light-dependent activity of Pba on enveloped viruses and suggest that the lipid membrane is the most likely target of the compound.

**Fig. 7.**
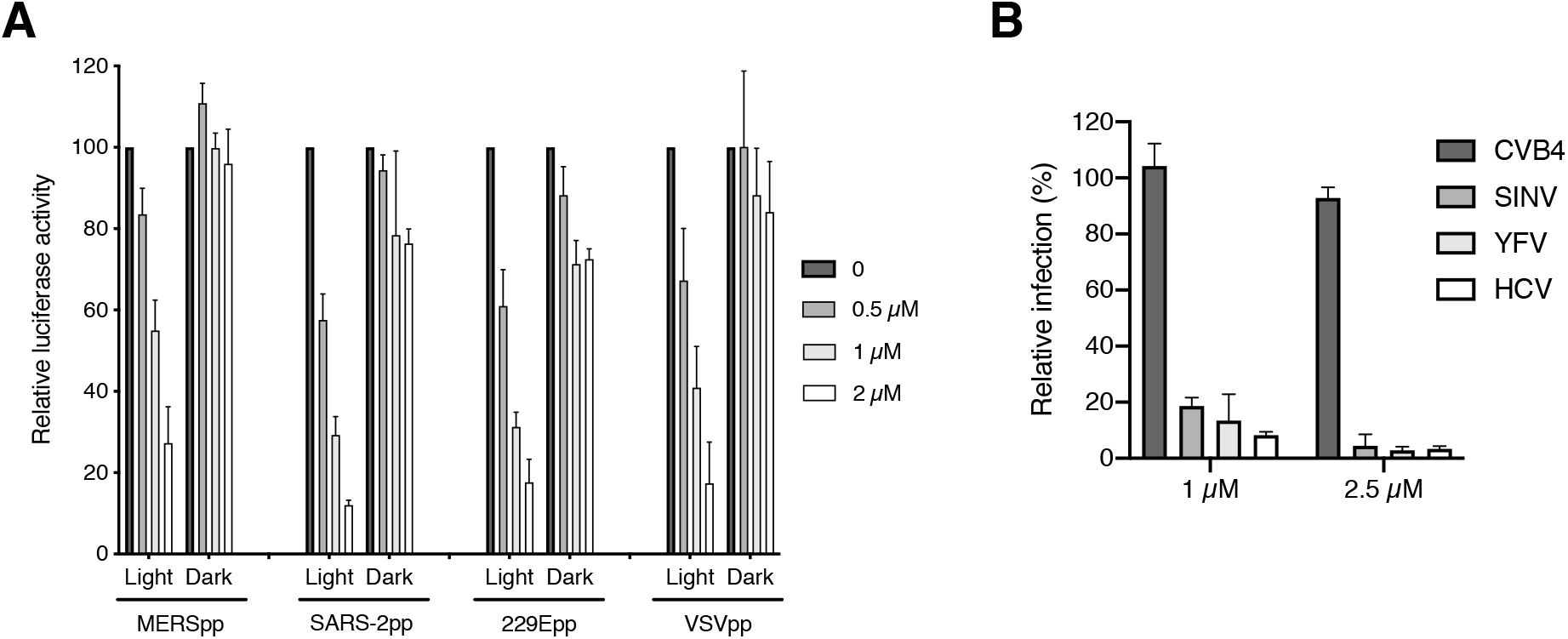
Pba is a broad-spectrum antiviral agent against enveloped viruses. (**A**) MERSpp, HCoV-229Epp, VSVpp, and SARS-2pp were preincubated with Pba at the indicated concentration either under light for 30 min (Light) or without light (Dark) prior to inoculation of Huh-7 cells expressing ACE2 and TMPRSS2 for 2 h. At 46 h post-inoculation, cells were lysed and luciferase activity was quantified. Infectivity is expressed as the percentage relative to the control (DMSO) to which the 100% value was arbitrarily attributed. Mean values ± SEM (error bars) of three different experiments are presented. (**B**). Different viruses were incubated with Pba at 1 and 2.5 μM under light condition for 30 min prior to inoculation. Cells were fixed at different time points depending on the virus (see Materials and Methods section for details) and subjected to immunofluorescence labelling. Infectivity is expressed as the percentage relative to the control (DMSO). Mean values ± SEM (error bars) of three different experiments are presented.

### Other chlorophyll-derived products and photosensitizers possess a light-dependent anti-coronaviral activity

Pba is a break-down product of chlorophyll. Chlorophyll is metabolized into different compounds including Pba, pyropheophorbide a (pyroPba), and chlorin e6. We wondered if these products would also have antiviral activity. Furthermore, we selected nine porphyrins or metalloporphyrins structurally related to Pba (N-methyl protoporphyrin IX, N-methyl mesoporphyrin IX, Zn-protoporphyrin IX, tin-mesoporphyrin IX, temoporfin, phthalocyanine, hemin chloride, HPPH, and 5,15-DPP), and one photosensitizer without related structure (Rose Bengale) to determine if similar antiviral activity against coronaviruses could be identified. The toxicity and antiviral activity of these compounds were investigated (Fig. S3). Chlorophyll b, phthalocyanine and 5,15-DPP were not active at the tested concentrations. Three molecules, hemin chloride, temoporfin and Rose Bengale had a moderate antiviral activity. The antiviral activity of the six most active compounds was tested both under normal white light conditions and in the dark, clearly showing that, similar to Pba, the antiviral activity of most molecules tested was also light-dependent (Fig. S3). Interestingly, N-methyl protoporphyrin IX and N-methyl mesoporphyrin IX showed an antiviral activity in dark conditions, but much lower than under light exposure. PyroPba was toxic at tested concentrations, thus dose-response experiments with lower concentrations were performed to determine precise CC_50_ and IC_50_. This was also done for all the active compounds. The CC_50_ and IC_50_ of the different compounds were compared and only pyroPba was more active against HCoV-229E than Pba (Table 1) with an IC_50_ of 0.35 μM. However, this compound is also more toxic with a CC_50_ of 2.67 μM and a selective index of 7.6. Thus, our results show that porphyrin-related compounds have antiviral activity against HCoV-229E but that Pba and pyroPba are the most active under normal white light exposure.

**Table 1.** Inhibitory and cytotoxic concentrations, and selective index of photosensitizers against HCoV-229E.

| **Molecules** | **IC<sub>50</sub> (μM)** | **CC<sub>50</sub> (μM) 24 h** | **SI** |
| --- | --- | --- | --- |
| Pba | 0.54 | 6.08 | 11.2 |
| PyroPba | 0.35 | 2.67 | 7.6 |
| Chlorin e6 | 0.72 | 21.4 | 29.7 |
| HPPH | 1.14 | 8.13 | 7.1 |
| N-methyl protoporphyrin IX | 1.21 | ND > 80 | >66.1 |
| N-methyl mesoporphyrin IX | 1.25 | ND > 80 | >64 |
| Zn protoporphyrin IX | 0.79 | 16.64 | 21.0 |
| Rose Bengale | 2.86 | ND > 80 | 27.9 |

### Pba reduces SARS-CoV-2 and MERS-CoV replication in human primary airway epithelial cells

As shown above, Pba is able to inhibit SARS-CoV-2 and MERS-CoV infection under white light exposure in cell culture. To determine if Pba could be used *in vivo*, its antiviral activity was tested in a preclinical model, the human primary airway epithelial cells. These cells, Mucilair™, are primary bronchial epithelial cells reconstituted in a 3D structure to mimic bronchial epithelium with an air-liquid interface. Mucilair™ cells were inoculated with SARS-CoV-2 or MERS-CoV in the presence of Pba at 0.25 or 2.5 μM. Remdesivir at 5 μM was used as a positive control. At 72 h post-inoculation, viral titers were determined and viral RNA levels were quantified. Viral RNA levels of SARS-CoV-2 and MERS-CoV were significantly decreased in cells in the presence of Pba at 0.25 and 2.5 μM, respectively (Fig. 8A). Similarly, viral titers of both viruses were decreased of more than 1xLog10 in the presence of Pba at 2.5 μM (Fig. 8B). These results confirm the antiviral activity of Pba against highly pathogenic human CoVs and its potential activity *in vivo*.

**Fig. 8.**
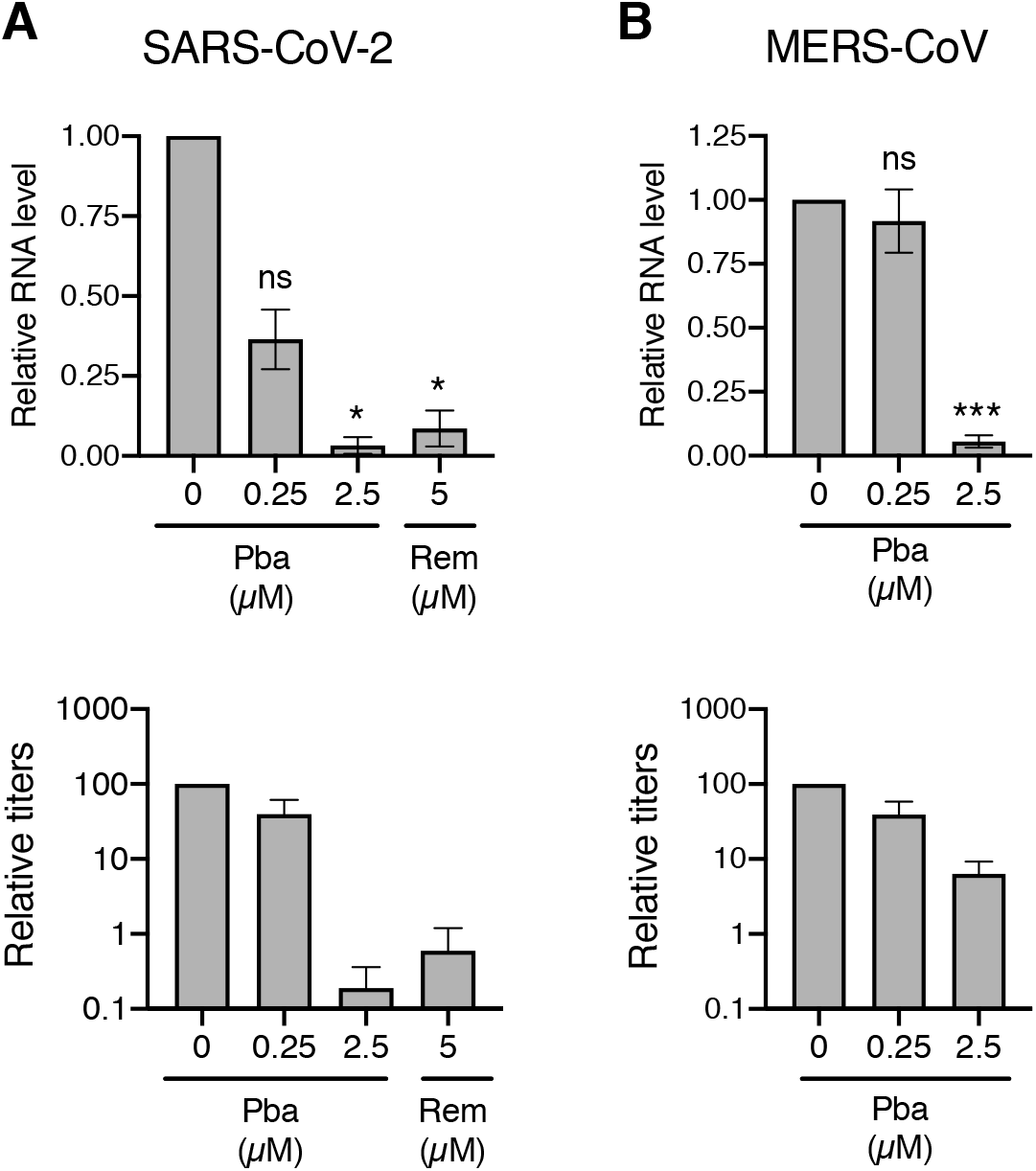
Antiviral efficacy of Pba in human primary bronchial epithelial cells. Mucilair™ cells were inoculated with SARS-CoV-2 or MERS-CoV in the presence of Pba at 0.25 or 2.5 μM for 1 h at the apical side. Inoculum was removed. Remdesivir (Rem) at 5 μM was added in the basolateral medium. 72 h post-inoculation, viruses were collected from the apical surface, and cells were lysed to extract RNA. Viral RNA was quantified by qRT-PCR and viral titers were determined by infectivity titrations for SARS-CoV-2 (**A**) and MERS-CoV (**B**). For RNA quantification, data are expressed relative to the control DMSO. Results are expressed as mean ± SEM of 3 experiments. Viral titers are representative of three independent experiments. *, P<0.05; ***, P<0.005; n.s., not significant.

## Discussion

By screening plant extracts for their antiviral activity against coronaviruses, the present study identified Pba as the active antiviral compound in the crude methanol extract from *M. opositifolius* after bioguided fractionation. It was demonstrated that Pba is active against various human CoVs, including SARS-CoV-2, HCoV-229E and MERS-CoV, as well as other enveloped viruses, including HCV, SINV and YFV, and various pseudotyped particles. Furthermore, we characterized Pba as a broad-spectrum antiviral photosensitizer causing PDI of all tested enveloped viruses by production of singlet oxygen species that most probably increase the rigidity of the lipid bilayer of the viral envelope.

Pba is a product of chlorophyll breakdown which is abundantly present in various plants (such as spinach) and marine algae. In general, the chlorophyll content in plants may vary and depends on the season, the part of the plant, the maturity of the organs and many other factors (*29*, *30*). Chlorophyll is degraded into Pba by chlorophyllase and some plants with high chlorophyllase content may contain more Pba (*31*, *32*). Pba is well documented for its potential as anti-cancer agent in photodynamic therapy (PDT). It is known to have a low toxicity, to selectively accumulate in tumours and to have a high adsorption at 665 nm (*33*, *34*).

For many years, photosensitizers have mainly been used as antitumor therapy, for which many photosensitizers have already been proven to be clinically safe and some are currently approved for use in humans (*35*). Although reports on the PDI of viruses go back to 1960, where it was shown that some photosensitive dyes, such as methylene blue, had an antiviral effect, it is only in the last decades that photosensitizers have gained considerable interest as antimicrobial (bacteria, fungi, and viruses) agents, due to their strong antimicrobial effects and low toxicity in normal tissue (*36*, *37*). A major advantage of those molecules is that, due to their direct damaging effect on the micro-organism, they are insensitive to the onset of resistance of the microorganism against the compound, the latter being a major problem in today’s research on antivirals and antibiotics. With the present study, we add Pba to the list of photosensitizers with considerable antiviral effects, at least against enveloped viruses. This antiviral effect is not new, since the antiviral activity of Pba has already been demonstrated before against several enveloped viruses, including HCV, Influenza A, herpes simplex-2 virus, and HIV-1 (*26*–*28*). Some studies also showed the direct effect on the viral particle, which is in line with our results. However, none of those reports showed any dependency on light exposure, and hence did not show that the antiviral effect is mediated by PDI of the particle. It is interesting to note that Pba (or highly related compounds such as pyroPba) has been isolated from different plant species and different organisms including marine algae (*38*), rendering Pba a very attractive antiviral due to its high availability.

Here, we show that Pba inhibits virus-cell fusion, probably by targeting and photodynamically damaging the viral membrane. With the help of cryo-EM, we demonstrated that the treatment of virions with Pba and exposed to the light did not affect their shape despite an osmotic shock. This is an indirect demonstration of an increased rigidity of the viral envelope upon Pba treatment. This feature was already demonstrated using a biophysical approach with lipophilic photosensitizers with antiviral activity (*24*, *39*). It was postulated that the increased rigidity impairs membrane bending required for viral fusion (*40*). Contrary to the cell membrane, the viral envelope is not able to undergo regeneration, which renders the PDI virus specific and insensitive to the onset of resistance. In the present study, several other compounds structurally related to Pba and to other known photosensitizers were also screened for their anti-coronaviral activity in order to find out whether other compounds would have a more potent effect. Several of those compounds, including pyroPba, chlorin e6, HPPH, N-methyl protoporphyrin IX, N-methyl mesoporphyrin IX, and Zn-protoporphyrin IX had a light-dependent antiviral effect, but only pyroPba turned out to be more active than Pba. However, pyroPba was also more toxic, and hence the final selectivity index was not higher than that of Pba. PyroPba has already been demonstrated to be active against IAV, RSV (respiratory syncytial virus), and SARS-CoV-2 **(*41*)**. The authors studied the mechanism of action of pyroPba against IAV and showed that the molecule targets the membrane of the virus and not the surface glycoproteins, a mechanism which is consistent with the one that we observed for Pba in our study. A requirement for photo-activation of pyroPba was not investigated nor mentioned by Chen *et al.* **(*41*)**. Porphyrins have already been described for their antiviral activity **(*25***, ***42*)**. Interestingly, three photosensitizers which have been shown to be active against VSV, including N-methyl protoporphyrin IX, N-methyl mesoporphyrin IX, and Zn-protoporphyrin IX, were also identified in our screen **(*43*)**. The authors clearly demonstrated that these compounds inactivated VSV after photoactivation via singlet oxygen release. We did not demonstrate the mechanism of action of these three molecules against HCoV-229E but we also demonstrated that they are active after photoactivation. Very recently, protoporphyrin IX and verteporfin were identified as inhibitors of SARS-CoV-2 **(*44***, ***45*)**. Both studies showed that protoporphyrin IX is active at an early step of infection, probably the entry step. Gu *et al.* postulated that the interaction of the compounds with ACE2 might impair the interaction of the virus with its receptor **(*44*)**. Lu *et al.* showed that protoporphyrin IX is active against several enveloped viruses but that the activity of protoporphyrin IX against IAV is not dependent on light activation **(*45*)**. Chlorin e6 is one of the most active compounds against HCoV-229E identified in this study. The antiviral activity of chlorin e6 against enveloped viruses such as HBV (Hepatitis B virus), HCV, HIV, DENV (dengue virus), MARV (Marburg virus), TCRV (tacaribe virus) and JUNV (Junin virus) has been already demonstrated **(*46*)**. Interestingly the authors also showed that the molecule is inactive against non-enveloped viruses, suggesting that it targets the viral envelope.

As mentioned above, many other photosensitizers have been studied for their antiviral activity (*25*, *40*), and for some of them, the PDI was clearly demonstrated as the mechanism of action (*40*, *47*). In light of the current SARS-CoV-2 pandemic, photosensitizers have received renewed attention as antiviral strategies to face this pandemic, and the use of those substances for the treatment of COVID-19 or the inactivation of SARS-CoV-2 on surfaces or in water has been postulated (*48*, *49*). Pba might have some advantages above the already described photosensitizers, as it is a highly available natural product and active under normal light conditions. Importantly, it does not require a very specific wavelength-dependent illumination treatment, at least not when applied on surfaces/mucosae exposed to the environmental light. However, the light-dependency of such molecules might render their application as therapeutic agents for internal organs (such as lungs for SARS-CoV-2) more challenging. Indeed, additional illumination will make its application as real therapy more complex, though not impossible because PDT is already used for the treatment of lung cancer (*50*). Efforts should be made for the development of specific device allowing PDT for COVID patients. Nonetheless, we believe that broad-spectrum, low-toxic, non-resistance inducing molecules such as Pba can certainly prove their value to reduce environment-to-person and person-to-person transmission of microorganisms when applied as e.g a spray for decontamination of surfaces or when formulated for topical application in nose and mouth. Very recently, a study describing such topical application of synthetic SARS-CoV-2 fusion inhibitor has demonstrated that the topical treatment of upper respiratory tract infections might prove its value in reducing virus transmission, particularly in cases where many people gather (*51*). In contrast to the SARS-CoV-2 fusion inhibitor described by de Vries *et al.* (*51*), Pba is a widely commercially available natural molecule with broad-spectrum activity against many enveloped viruses. Therefore, one should explore whether it can exert similar effects upon topical administration to the nose or oral cavity. If so, Pba might help to make people less susceptible to and/or less contagious upon upper respiratory infections with enveloped viruses, many of them causing seasonal outbreaks of respiratory disease such as common colds and flu.

Given that 1) onset of resistance to this product is very unlikely, 2) the activity of the compound is not dependent of envelope variants, 3) coronaviruses and other enveloped viruses can cause major problems in animals and 4) there is a potential risk for virus transmission from those animal to humans, the possibility to formulate Pba in such a way that also veterinary medicine and facilities with large numbers of animals can benefit from the strong antiviral properties that this molecule might have in the environment (decontamination of air, water and surfaces) should additionally be explored.

## Materials and Methods

### Chemicals

Dulbecco’s modified Eagle’s medium (DMEM), Opti-MEM, phosphate buffered saline (PBS), 4’,6-diamidino-2-phenylindole (DAPI), were purchased from Life Technologies. Goat and foetal bovine sera (FBS) were obtained from Eurobio. Pheophorbide a (Pba) >90% pure, pyroPba, chlorin e6, HPPH, N-methyl protoporphyrin IX, N-methyl mesoporphyrin IX, and Zn protoporphyrin IX were from Cayman chemicals (Merck Chemicals, Darmstadt, Germany). Remdesivir (GS-5734) was from Selleck Chemicals (Houston TX). Mowiol 4-88 was obtained from Calbiochem. Rose Bengale, trolox and other chemicals were from Sigma (St. Louis, MO). Stocks of compounds were resuspended in dimethylsulfoxide (DMSO) at 50 mM. Plant extracts were resuspended in DMSO at 25 mg/mL.

### Antibodies

Mouse anti-HCV E1 mAb A4 (*52*) and mouse anti-YFV E mAb 2D12 (anti-E, ATCC CRL-1689) were produced *in vitro* by using a MiniPerm apparatus (Heraeus). Mouse anti-dsRNA mAb (clone J2) was obtained from Scicons. Mouse anti-SARS-CoV-2 spike protein mAb were obtained from GeneTex. Polyclonal rabbit anti-SARS-CoV-2 nucleocapsid antibodies were from Novus. Cyanine 3-conjugated goat anti-mouse IgG and HRP-labeled goat-anti rabbit IgG antibodies were from Jackson Immunoresearch.

### Cells and culture conditions

Huh-7, Vero-81 (ATCC number CCL-81) and Vero-E6 were grown in DMEM with glutaMAX-I and 10% FBS in an incubator at 37°C with 5% CO_2_. Vero-81 cells were subcloned to obtain a better overall infection rate. The primary human bronchial epithelial cells Mucilair™ were from Epithelix (Geneva, Switzerland) and maintained in Mucilair™ culture medium (Epithelix) as recommended by the manufacturer.

### Plant collection and extraction

The fifteen plants were collected in the Bafing region (North-West Côte d’Ivoire, Touba department). They were authenticated at the Centre National de Floristique (CNF), University of Félix Houphouët Boigny de Cocody (Abidjan), where voucher specimens were deposited in an Herbarium. *M. oppositifolius* voucher number is UCJ006172. Plants were cleaned and air-dried at constant temperature (26°C) for 1 to 2 weeks at the Nangui Abrogoua University (Abidjan). They were then powdered and stored in the dark until extractions. For each plant, 20 g of dried powder were mixed with 100 mL methanol for 24 h. After filtration, the grounds were extracted again twice in the same way. The 3 resulting filtrates were combined and dried under vacuum at 40°C. These extracts were then dissolved in DMSO for antiviral assays.

### Bioguided fractionation of Mo extract and Pba identification

For *Mallotus oppositifolius*, three other solvents were used to extract more compounds from these plant leave: methylene chloride (MC) for the first extraction of the dried leaves, then methanol to extract the first ground, and ethanol/water (50:50) to extract the second ground. The corresponding extracts were tested against HCoV-229E-Luc. Since the MC extract was the most active, it was fractionated by chromatography (CPC), leading to 10 fractions (F1-10) that were tested again. F7 was selected for further fractionation by another chromatography (HPLC) and led to 9 partitions (7.1-7.9). Partition 7.7 (the most active against HCoV-229E) purity and identity was determined by UPLC-MS and NMR (more details in supplementary materials).

### Viruses

The following viral strains were used: HCoV-229E strain VR-740 (ATCC), and a recombinant HCoV-229E-Luc (kind gift of Pr. V. Thiel) (*53*); SARS-CoV-2 (isolate SARS-CoV-2/human/FRA/Lille_Vero-81-TMPRSS2/2020, NCBI MW575140) was propagated on Vero-81-TMPRSS2 cells. MERS-CoV was recovered by transfecting the infectious clone of MERS-CoV-EMC12 (kindly provided by Dr Luis Enjuanes) in Huh-7 cells. A cell culture-adapted strain (JFH1-CSN6A4) of HCV was produced as previously described (*54*). A recombinant Sindbis virus (SINV) expressing HCV E1 glycoprotein was employed as previously described (*55*). Yellow fever virus strain 17D (YFV) was obtained from Dr Philippe Desprès (Institut Pasteur de Paris, France). Coxsackievirus B4 strain E2 (CVB4) was provided by Dr Didier Hober (Université de Lille, France).

### HCoV-229E infection inhibition assays

#### Luciferase assay

HCoV-229E-Luc was first mixed with the crude extracts or the compounds at the appropriate concentrations for 10 minutes. Huh-7 cells and Huh-7-TMPRSS2 cells were inoculated with HCoV-229E-Luc at a MOI of 0.5 in a final volume of 50 μL for 1 h at 37°C in the presence of the plant crude extracts or the different compounds. The virus was removed and replaced with culture medium containing the extracts or the different compounds for 6 h at 37°C. Cells were lysed in 20 μL of Renilla Lysis Buffer (Promega, Madison, USA) and luciferase activity was quantified in a Tristar LB 941 luminometer (Berthold Technologies, Bad Wildbad, Germany) using Renilla Luciferase Assay System (Promega) as recommended by the manufacturer.

This experiment was either performed under white light exposure, in which the virus and the compounds were exposed to the light of the BSC lamp. To maximize light exposure, the tubes were laid flat on the bench of the BSC. For dark condition, the light of the BSC and the room was shut down and all the tubes and plates were covered with foil paper.

#### HCoV-229E titers

Huh-7 and Huh-7-TMPRSS2 cells seeded in 24-well plates were inoculated with HCoV-229E at a MOI of 0.5 in the presence of Pba at different concentrations for 1 h at 37°C. The inoculum was removed and replaced with culture medium containing Pba and the cells were incubated at 37°C for 8 h (for TMPRSS2 condition) or 10 h (without TMPRSS2). Supernatants were collected and serial dilutions were performed and used to infect naïve Huh-7 cells in 96-well plates. Six days after infection, cytopathic effect was determined in each well to calculate TCID_50_ titers by using the Reed and Muench method.

### SARS-CoV-2 and MERS-CoV infection inhibition assays

Vero-E6 and Huh-7 cells seeded in 24-well plates 24 h before inoculation were inoculated with SARS-CoV-2 and MERS-CoV, respectively, at a MOI of 0.3 in the presence of Pba at different concentrations for 1 h at 37°C. Then, the inoculum was removed by 3 washings with DMEM and fresh medium containing different Pba concentrations was added for 16 h at 37°C. Cell supernatants were collected and the amount of infectious virus was determined by infectivity titration. Therefore, Vero-E6 (SARS-CoV-2) and Huh-7 (MERS-CoV), seeded in 96-well plates, were inoculated with 100 μL of 1/10 serially diluted supernatants (ranging from 10^−1^ to 10^−8^). Cells were incubated with the virus dilutions for 5 days at 37°C and 5% CO_2_. Then, the 50% tissue culture infectious dose (TCID_50_) was determined by assessing the CPE in each well by light microscopy and the 50% end point was calculated according to the method of Reed and Muench.

### Time-of-addition assay

To determine at which stage of the replication cycle Pba executed its effect, a time-of-addition assay was performed for which 1μM Pba (and 10 μM chloroquine as a control for SARS-CoV-2) was added at different time points before (referred to as the condition ‘pre-treatment cells’), during (referred to as the condition ‘inoculation’), or after inoculation. For the latter condition, Pba and chloroquine were not added before and during inoculation, but only directly after removal of the inoculum (referred to as the condition ‘p.i. - end’), or from 1 h or 2 h after removal of the inoculum onwards (referred to as the condition ‘1 h p.i.-end’ and ‘2 h p.i.-end, respectively) and were left in the medium for the rest of the incubation time (i.e until 6 h p.i. for HCoV-229E and 16 h p.i. for SARS-CoV-2). For this experiment, Huh-7-TMPRSS2 or Vero-81 cells were inoculated with HCoV-229E-Luc or SARS-CoV-2 at a MOI of 0.5 and 0.05, respectively. One hour after inoculation, cells for all conditions were washed 3 times to remove the unbound particles. HCoV-229E-luc, luciferase activity was quantified as described ahead. For SARS-CoV-2, cells were washed once with PBS and lysed in 200 μL of non-reducing 2x Laemmli loading buffer. Lysates were incubated at 95°C for 30 min to inactivate the virus and lysates were kept at −20°C until western blot analysis (see below). For each time point, DMSO was taken as a control, and all experiments were repeated 3 times.

### Western blot detection of the SARS-CoV-2 nucleocapsid expression

Sixteen hours after inoculation, cells were washed once with PBS and lysed in 200 μL of non-reducing 2x Laemmli loading buffer. Lysates were incubated at 95°C for 30 min to inactivate the virus and the proteins were subsequently separated on a 12% polyacrylamide gel by SDS-PAGE. Next, proteins were transferred to a nitrocellulose membrane (Amersham), and the membranes were subsequently blocked for 1 h at RT in 5% (w/v) non-fat dry milk in PBS with 0.1% (v/v) Tween-20. Membranes were incubated overnight at 4°C with polyclonal rabbit anti-SARS-CoV-2 nucleocapsid antibodies in 5% (w/v) non-fat dry milk in PBS with 0.1% (v/v) Tween-20. After being washed 3 times with PBS with 0.1% (v/v) Tween-20, membranes were incubated for 1 h at RT with HRP-labeled goat-anti rabbit IgG antibodies, after which membranes were washed 3 times. N proteins were visualized by enhanced chemiluminescence (Pierce™ ECL, ThermoFisher Scientific). Quantification was performed by using Image J and its gel quantification function.

### Infection assay with other viruses

Vero (YFV, SINV, CBV4) or Huh-7 (HCV) cells grown on glass coverslips were infected with viral stocks diluted so as to obtain 20–40% infected cells in control conditions. The cells were fixed at a time that allowed for a clear detection of infected cells vs non-infected cells, and avoided the detection of reinfection events, thus limiting the analysis to a single round of infection (30 h p.i. for HCV, 20 h p.i. for YFV, 6 h p.i. for SINV, 4 h p.i. for CVB4). The cells were fixed for 20 min with 3% PFA. They were then rinsed with PBS and processed for immunofluorescence as previously described (*56*) using primary mouse antibodies specific to HCV E1 (for both HCV and SINV), YFV E, or dsRNA (for CVB4), followed by a cyanine-3-conjugated goat anti-mouse IgG secondary antibody for the detection of infected cells. Nuclei were stained with DAPI. Coverslips were mounted on microscope slides in Mowiol 4-88-containing medium. Images were acquired on an Evos M5000 imaging system (Thermo Fisher Scientic) equipped with light cubes for DAPI, and RFP, and a 10× objective. For each coverslip, a series of six 8-bit images of randomly picked areas were recorded. Cells labelled with anti-virus mAbs were counted as infected cells. The total number of cells was obtained from DAPI-labelled nuclei. Infected cells and nuclei were automatically counted using macros written in ImageJ. Infections were scored as the ratio of infected over total cells. The data are presented as the percentage of infection relative to the control condition.

### Effect of Pba on pseudotyped virion entry

Particles pseudotyped with either SARS-CoV-2 S (SARS-2pp), MERS-CoV S proteins (MERSpp), HCoV-229E-S (HCoV-229Epp), genotype 2a HCV envelope proteins (HCVpp), or the G envelope glycoprotein of vesicular stomatitis virus (VSV-Gpp) were produced as previously described (*22*, *57*). Pseudotyped virions were pre-treated with Pba for 30 min at room temperature under the BSC’s light or covered in foil and then used to inoculate Huh-7 cells in 96-well plates for 3 h. The inoculum was removed and cells were further incubated with culture medium for 45 h. Cells were lysed and luciferase activity was detected by using a Luciferase Assay kit (Promega) and light emission measured by using a Tristar LB 941 luminometer (Berthold Technologies).

### White light exposure kinetics

HCoV-229E-Luc or SARS-CoV-2 were pre-treated with Pba at room temperature and exposed to BSC’s white fluorescent light during different period of time. To maximize light exposure, tubes were laid flat under the BSC’s light. Next, infection was quantified for each virus as described previously.

### Fusion assay

Cells were preincubated for 30 min in the presence of 25 mM NH_4_Cl at 37°C to inhibit virus entry through the endosomal route, and then were transferred to ice. In the meantime, the virus was preincubated under light with Pba and 25 mM NH_4_Cl for 10 min and then allowed to bind to the cells at 4°C for 1 h in DMEM containing 0.2% BSA, 20 mM Hepes, and 25 mM NH_4_Cl. Cells were then warmed by addition of DMEM containing 3 μg/mL trypsin, 0.2% BSA, 20 mM Hepes, and 25 mM NH_4_Cl and were incubated for 5 min in a water bath at 37°C. The cells were rinsed and further incubated for 30 min in culture medium containing 25 mM NH_4_Cl, and then the medium was replaced by normal culture medium. Seven hours after inoculation, luciferase activity was detected by using a Renilla Luciferase Assay Kit (Promega).

### Cell-cell fusion assay by transient expression of the SARS-CoV-2 spike protein

Vero-81 cells were seeded on coverslips in 24-wells 16 h before transfection. Cells were transfected with 250 ng of a pCDNA3.1(+) vector encoding for the SARS-CoV-2 spike protein using the TransIT^®^-LT1 Transfection Reagent (Mirus Bio). Six hours post transfection (p.t.), transfection medium was replaced by normal medium containing 1 μM Pba or DMSO. Twenty-four hours p.t., cells were fixed with 3% paraformaldehyde in PBS for 20 min at RT and syncytia were visualized by immunofluorescence, by incubating the cells with a monoclonal anti-SARS-CoV-2-spike antibody in 10% normal goat serum, followed by incubation with Cyanine-3-conjugated goat anti-mouse IgG antibodies. Nuclei were visualized with 1 μg/ml of 4’,6-diamidino-2-phenylindole (DAPI), and coverslips were mounted in Mowiol^®^ mounting medium. Pictures were obtained with an Evos M5000 imaging system (Thermo Fisher Scientific).

### Attachment assay

Huh-7-TMPRSS2 cells seeded in 24-well plates were inoculated with HCoV-229E at a MOI of 4 on ice in the presence of 4.1 or 8.2 μM Pba under the light of the BSC. 1 h after inoculation, cells were washed 3 times with cold PBS, and lysed using LBP lysis buffer for RNA extraction following manufacturer’s instructions (NucleoSpin^®^ RNA plus extraction kit, Macherey-Nagel). Reverse transcription was then performed on 10 μL of RNA using High Capacity cDNA Reverse Transcription kit (Applied Biosystems). 3 μL of cDNA were used for real-time reverse-transcription polymerase chain reaction (qRT-PCR) assay using specific primers and probe targeting the N gene (forward primer 5′-TTCCGACGTGCTCGAACTTT-3′, reverse primer 5′-CCAACACGGTTGTGACAGTGA-3′ and probe 5′-6FAM-TCCTGAGGTCAATGCA-3’) and subjected to qPCR amplification with Taqman Master mix.

#### Quencher Assay

HCoV-229E-Luc was mixed with 10 mM Trolox or NaN_3_ after which 0.5 or 1 μM Pba was added and the mixture was exposed to light for 10 min. The mixture was used to inoculate Huh-7-TMPRSS2 cells for 1h. Inoculum was replaced with DMEM and cells were kept in the dark at 37°C 5% CO_2_ for 7 h and then lysed to quantify luciferase activity as described above.

#### CryoEM

HCoV-229E was produced by inoculating a confluent Huh-7 T75 Flask at MOI of 0.008 in DMEM supplemented with 5% FBS and put at 33°C 5% CO_2_ for 5 days. Supernatant was harvested and treated with DMSO or 10 μM Pba, and further kept in the dark or exposed to light for 30 min. Then NaCl was added to a final concentration of 400 mM final to induce an osmotic shock. Viruses were fixed in 4% PFA. For cryo-EM experiments of the particles, lacey carbon formvar 300 mesh copper grids were used after a standard glow discharged procedure. Plunge freezing was realized using the EM-GP apparatus (Leica). Specimens were observed at −175 °C using a cryo holder (626, Gatan), with a ThermoFisher FEI Tecnai F20 electron microscope operating at 200 kV under low-dose conditions. Images were acquired with an Eagle 4k x 4k camera (ThermoFisher FEI).

#### Primary airway cell infection quantification

The air interface of Mucilair™ (Epithelix) was rinsed with 100 μL of medium for 10 min 3 times to remove mucosal secretion. The cells were then inoculated at the apical membrane with SARS-CoV-2 or MERS-CoV at a MOI of 0.2 in the presence of compounds for 1 h at 37°C. Inoculum was removed and the cells were rinsed with PBS. In parallel, compounds were added in the basolateral medium. 72 h post-infection, viruses secreted at the apical membrane were collected by adding 200 μL of medium in the apical chamber. Viral titers were determined as described above. In parallel, cells were lysed with lysis buffer from the kit NucleoSpin^®^ RNA Plus (Macherey Nagel), and total RNA extracted following manufacturer’s instructions, eluted in a final volume of 60 μL of H_2_O, and quantified.

For SARS-CoV-2, one-step qPCR assay was performed using 5 μL of RNA and Takyon Low rox one-step RT probe Mastermix (Eurogentec) and specific primers and probe targeting E gene, forward primer 5’-ACAGGTACGTTAATAGTTAATAGCGT-3’, reverse primer 5’-ATATTGCAGCAGTACGCACACA-3’ and probe FAM-ACACTAGCCATC-CTTACTGCGCTTCG-MGB.

For MERS-CoV and RPLP0 reference gene, 10 μM of RNA were used for cDNA synthesis using High Capacity cDNA Reverse Transcription kit (Applied Biosystems). 3 μL of cDNA were used for real-time reverse-transcription polymerase chain reaction (qRT-PCR) assay using specific probes. For MERS-CoV, the following primers and probe targeting N gene were used, forward primer 5′-GGGTGTACCTCTTAATGCCAATTC-3′, reverse primer 5′-TCTGTCCTGTCTCCGCCAAT-3′ and probe 5′-FAM-ACCCCTGCGCAAAATGCTGGG-MGBNFQ-3′ and subjected to qPCR amplification with Taqman Master mix. For RPLP0, Taqman gene expression assay (Life Technologies) was used according to the manufacturer instruction. SARS-CoV-2 E and MERS-CoV N gene expression were quantified relative to RPLP0 using ΔΔCt method. A value of 1 was arbitrary assigned to infected cells without compound.

### Statistical analysis and IC_50_ and CC_50_ calculation

Values were graphed and IC_50_ calculated by non-linear regression curve fitting with variable slopes constraining the top to 100% and the bottom to 0%, using GraphPad PRISM software. Kruskal Wallis nonparametric test followed by a Dunn’s multicomparison post hoc test with a confidence interval of 95% was used to identify individual difference between treatments. P values < 0.05 were considered as significantly different from the control.

## Acknowledgments

We thank Volker Thiel for providing HCoV-229E-RLuc, Philippe Desprès for YFV, Luis Enjuanes for MERS-CoV and Didier Hober for CVB4. We are also grateful to Robin Prath and Nicolas Vandenabele for their technical help in the BSL3 facility. Authors are grateful to the LARMN platform (University of Lille, France) and wish to thank N. Azaroual and V. Ultré for their help on NMR analysis.

## Author contributions

Conceptualization: KS
SBo Methodology: KS, LD, SBo, SBe, OL, SS
Investigation: TM, LD, MB, NF, KH, YR, MD
Supervision: KS, SBo, SS, JD, FHTB
Writing—original draft: KS, LD, SBo
Writing—review & editing: SBe, JD, OL, YR, SS

## Data and materials availability

All data are available in the main text or the supplementary materials.

## Supplementary Text

### Materials and methods

#### Bioguided fractionation of Mo extract

*Mallotus oppositifolius* was extracted with three solvents of increasing polarity. For each solvent, 5 mL/g of dried powder was used for 24h. Each extraction was repeated 3 times and the resulting extracts were combined and vacuum dried. First, the three dried leave was macerated in methylene chloride (MC), then the first ground was macerated in methanol, and finally ethanol/water (50:50) was used to extract the second ground. These 3 dried extracts were dissolved in DMSO and tested against HCoV-229E-Luc. The MC partition (most active) was fractionated by Centrifugal Partition Chromatography (CPC, Armen instruments^®^). CPC is a liquid/liquid chromatography based on the partition of a sample in a biphasic immiscible liquid system. The system consists of a rotor connected to two Shimadzu^®^-LC-20AP pumps, a CBM-20A controller, and a SPD-M20A diode array detector. The Arizona S system (heptane/ethyl acetate/methanol/water, 5:2:5:2) composed the mobile and stationary phases. Injections were carried out with 3 g of the MC extract solubilized in 50 mL of a mixture of mobile and stationary phases (5:5). The analysis took 60 min at 30 mL/min and 1200 rpm. The extrusion took 35 min at 50 mL/min with the same rotor speed. This method allowed to obtain 10 different fractions (F1-10) that were vacuum dried, dissolved in DMSO and then tested again on HCoV-229E-Luc. Fraction F7 was the most active and selected for further fractionation by another chromatography. We used a preparative HPLC system composed of the same pumps, controller and detector as our CPC system. The stationary phase was a Vision HT HL C18 (5μm, 250×10 mm) column (Grace). The mobile phase was a mixture of methanol and water with the following gradient: 50-100% (0– 15 min), and 100% methanol (15-30 min). F7 was dissolved in methanol and injected repeatedly (500 μL at 20 mg/mL). The flow rate was set at 3 mL/min. This process led to 9 partitions (7.1-7.9).

#### Structural elucidation of compound in F7.7

Partition 7.7 was the most active against HCoV-229E and further analysed by Ultra-High Performance Liquid Chromatography (UPLC-UV-MS) and Nuclear Magnetic Resonance (NMR). UPLC-UV-MS analysis were performed on an Acquity UPLC^®^H-Class system (Waters, Guyancourt, France) coupled with a Diode Array Detector (DAD) and a QDa ESI-Quadrupole Mass Separation, using an ACQUITY UPLC^®^ BEH C18 1.7μm (2.1×100mm) column (Waters, Milford MA). Gradient elution was performed with (A) 0.1% formic acid in water and (B) 0.1% formic acid in acetonitrile at a flowrate of 0.3 mL/min, as following: 30-90% (0-3 min), 90-100% (3-7 min) before returning to the initial conditions (30% B). Analytes were monitored using UV detection (190 to 790 nm) and MS-Scan from 100 to 1000 Da (both in positive and negative mode). All data were acquired and processed using Empower 3 software.

The structural elucidation of F7.7 was conducted with NMR. Monodimensional spectra (1H and 13C) were recorded on a Bruker DPX-500 spectrometer. The chemical structure was established by comparison with literature data (*1*).

#### Cell toxicity assay

6×10^4^ Huh-7, Vero-E6 and Vero-81 cells were seeded in 96-well plates and incubated for 16 h at 37°C 5% CO_2_ incubator. The cells were then treated with increasing concentrations of the compound of interest. One hour after inoculation, cells were either left in the incubator (dark condition) or taken out to be exposed to the white light of the biosafety cabinet (BSC) for 10 min (light condition), after which cells were further incubated in the dark at 37°C 5% CO_2_ for 23 h.

BSC’s light source lamp consists of one fluorescent tube of 36W, 3350 lumen white light. An MTS [3-(4,5-dimethylthiazol-2-yl)-5-(3-carboxymethoxyphenyl)-2-(4-sulfophenyl)-2H-tetrazolium]-based viability assay (Cell Titer 96 Aqueous non-radioactive cell proliferation assay, Promega) was performed as recommended by the manufacturer. The absorbance of formazan at 490 nm was detected using a plate reader (ELX 808 Bio-Tek Instruments Inc). Each measure was performed in triplicate and each experiment was repeated at least 3 times.

---

**Fig. S1.**
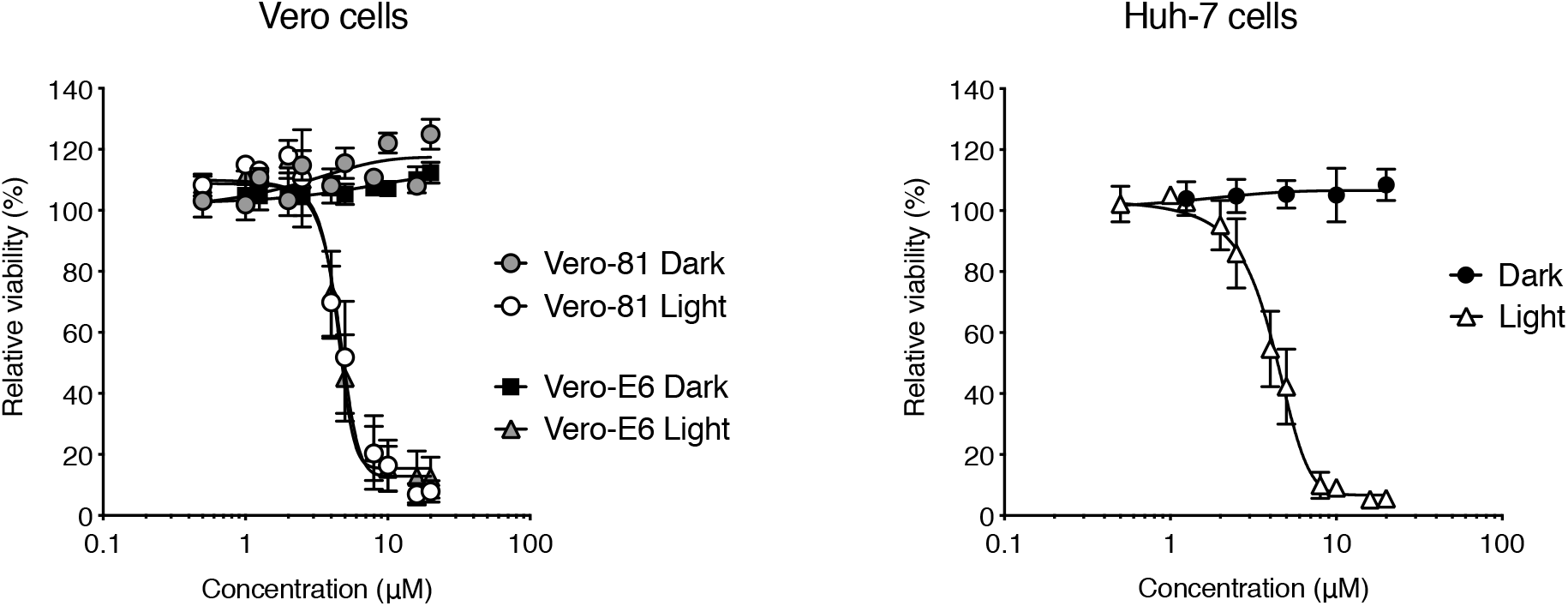
Toxicity of Pba depends on light exposure. Cells were incubated with Pba at different concentrations and either kept in the incubator for 24 h (Dark), or were taken out of the incubator after 1 h of incubation with Pba, and left for 10 min under light exposure, after which the cells were replaced in the incubator for 23 h (Light). Data are expressed relative to the control DMSO. Results are expressed as mean ± SEM of 3 experiments.

**Fig. S2.**
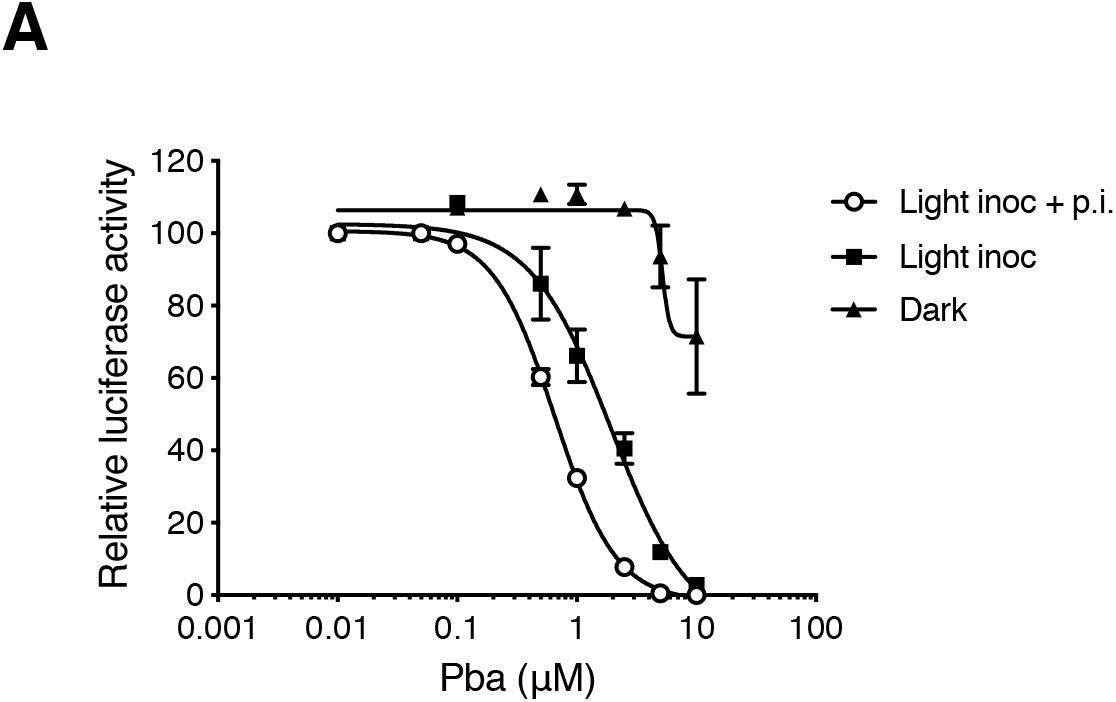
The antiviral activity of Pba is light-dependent. Huh-7 cells were inoculated with HCoV-229E-Luc in the presence of various concentrations of Pba either with the light of the laminar flow cabinet turned on (Light inoc) or off (Dark). One hour after inoculation, the inoculum was removed, either in light (Light inoc + p.i.) or dark conditions (Light inoc), and cells were further incubated with Pba for 6 h, after which luciferase activity was measured.

**Fig. S3.**
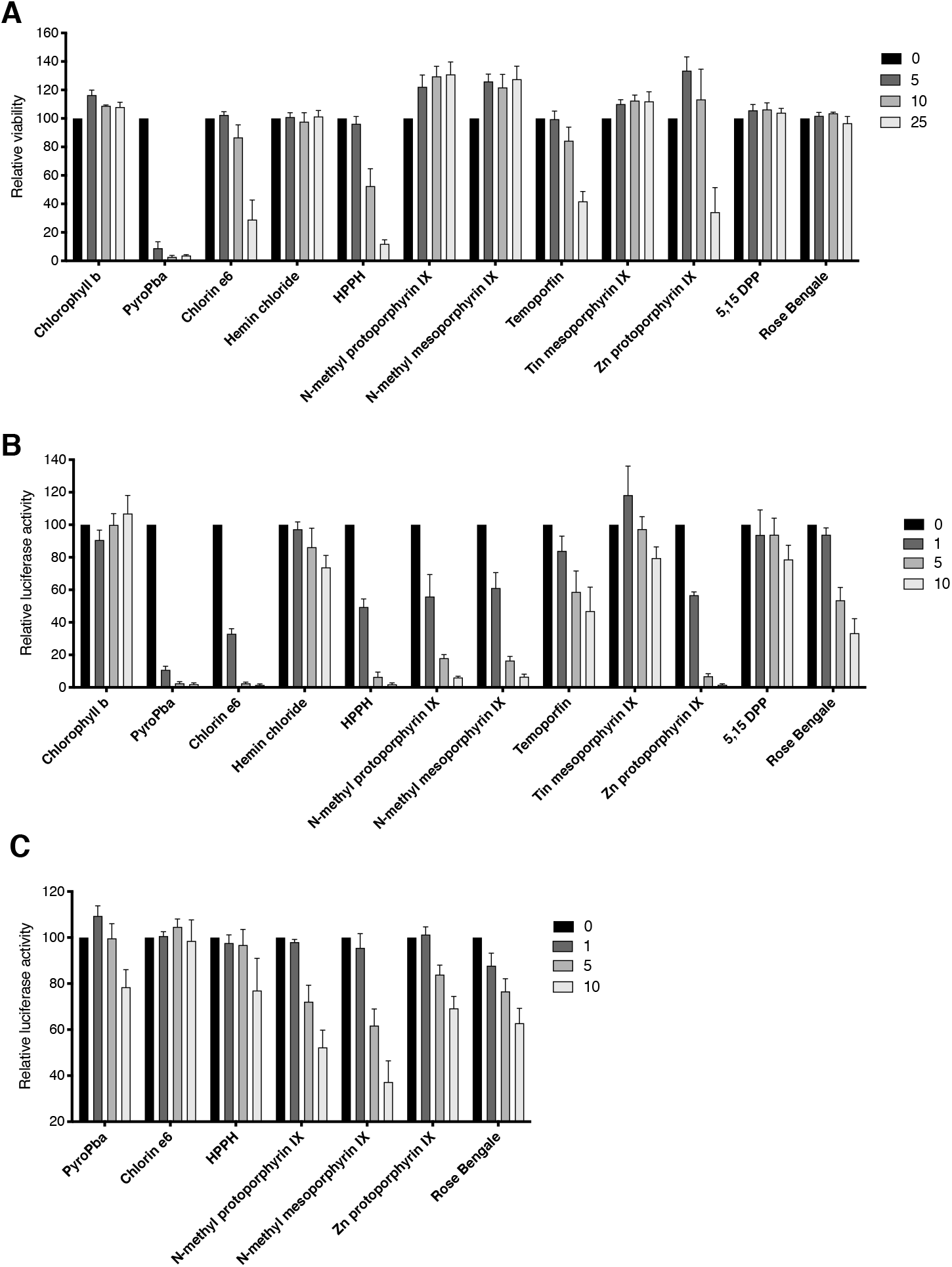
Activity of structurally-related Pba compounds and other photosensitizers on HCoV-229E infection. **A**. The toxicity on Huh-7 cells of the different compounds was determined by MTS assay. Huh-7 cells were incubated with the molecules at 5, 10, and 25 μM under the light of the cabinet. The medium was removed after 1 h and exposed for 10 min to the light of the cabinet to mimic infection assay, then placed in the dark for 23 h and MTS assay was performed. **B** and **C**. Huh-7 cells were inoculated with HCoV-229E-Luc in the presence of indicated compounds at different concentrations either under light exposure (**B**) or in the dark (**C**). 1 h post inoculation, the inoculum was removed and replace with fresh medium containing the compounds and exposed or not for 10 min to the light of the cabinet. Cells were lysed 7 h p.i. to quantify luciferase. Data are expressed relative to the control DMSO. Results are expressed as mean ± SEM of 3 experiments.

