## Supplementary data for "A photoactivable natural product with broad antiviral activity against enveloped viruses including highly pathogenic coronaviruses"

**Title**

- **A photoactivable natural product with broad antiviral activity against enveloped viruses including highly pathogenic coronaviruses**
- Short title: A new coronavirus fusion inhibitor

**Authors**

Thomas Meunier†[1], Lowiese Desmarets†[1], Simon Bordage†[2], Moussa Bamba[2,3], Kévin Hervouet[1], Yves Rouillé[1], Nathan François[1], Marion Decossas[4], Fézan Honora Tra Bi[3], Olivier Lambert[4], Jean Dubuisson[1], Sandrine Belouzard[1], Sevser Sahpaz[2]‡, and Karin Séron[1]‡*

**Fig. S1**

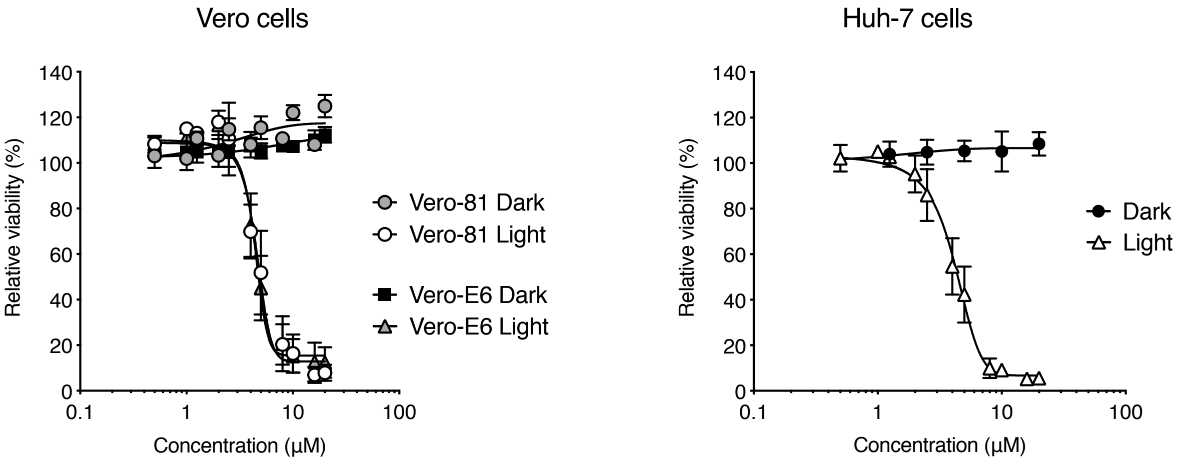


**Fig. S1. Toxicity of Pba depends on light exposure.** Cells were incubated with Pba at different concentrations and either kept in the incubator for 24 h (Dark), or were taken out of the incubator after 1 h of incubation with Pba, and left for 10 min under light exposure, after which the cells were replaced in the incubator for 23 h (Light). Data are expressed relative to the control DMSO. Results are expressed as mean ± SEM of 3 experiments.

**Fig. S2**

**A**

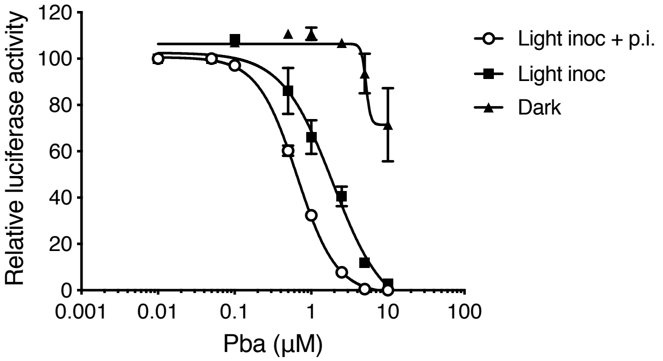


**Fig. S2. The antiviral activity of Pba is light-dependent.** Huh-7 cells were inoculated with HCoV-229E-Luc in the presence of various concentrations of Pba either with the light of the laminar flow cabinet turned on (Light inoc) or off (Dark). One hour after inoculation, the inoculum was removed, either in light (Light inoc + p.i.) or dark conditions (Light inoc), and cells were further incubated with Pba for 6 h, after which luciferase activity was measured.

**Fig. S3**

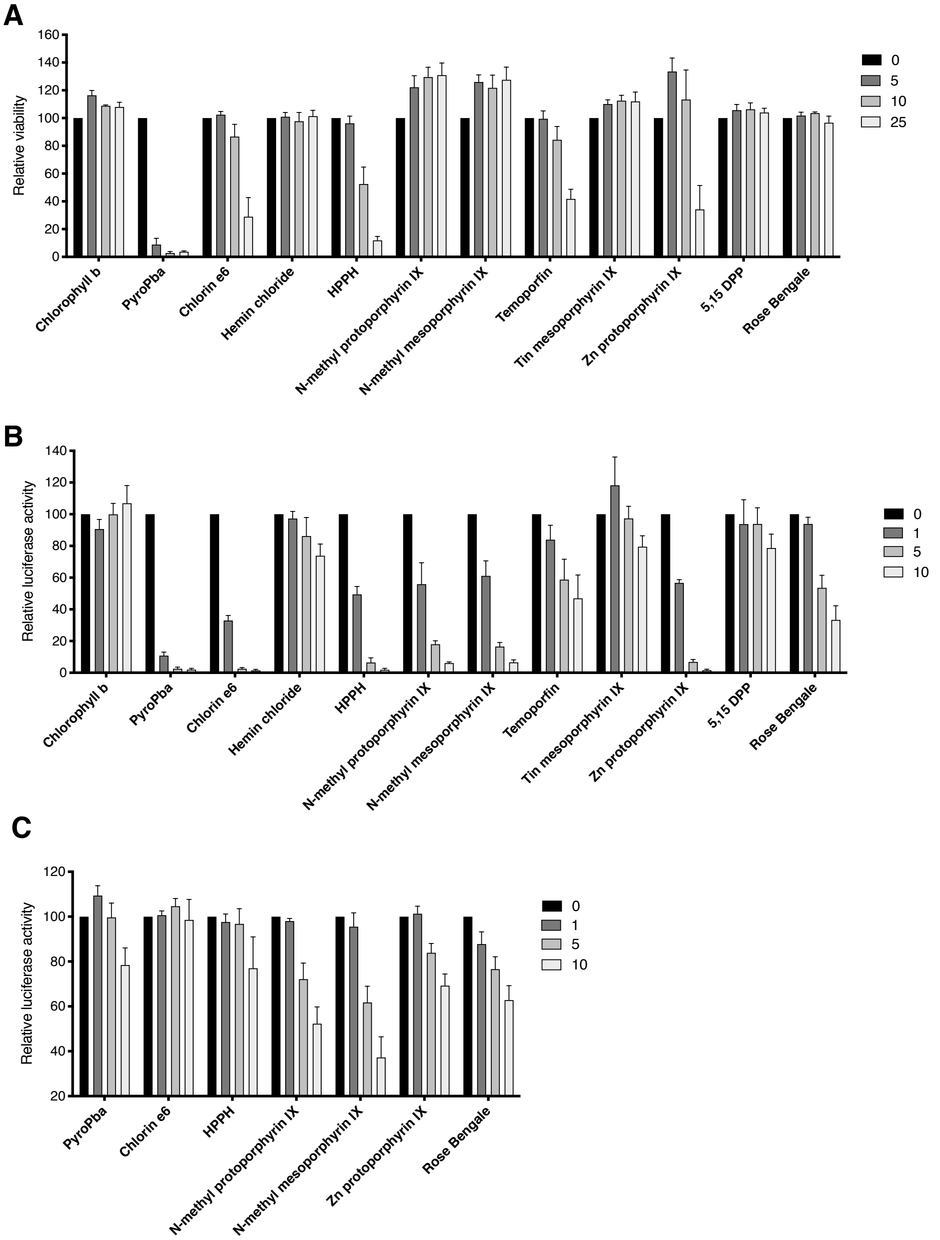


**Fig. S3. Activity of structurally-related Pba compounds and other photosensitizers on HCoV-229E infection. A.** The toxicity on Huh-7 cells of the different compounds was determined by MTS assay. Huh-7 cells were incubated with the molecules at 5, 10, and 25 µM under the light of the cabinet. The medium was removed after 1 h and exposed for 10 min to the light of the cabinet to mimic infection assay, then placed in the dark for 23 h and MTS assay was performed. **B** and **C**. Huh-7 cells were inoculated with HCoV-229E-Luc in the presence of indicated compounds at different concentrations either under light exposure (**B**) or in the dark (**C**). 1 h post inoculation, the inoculum was removed and replace with fresh medium containing the compounds and exposed or not for 10 min to the light of the cabinet. Cells were lysed 7 h p.i. to quantify luciferase. Data are expressed relative to the control DMSO. Results are expressed as mean ± SEM of 3 experiments.
